# LoGicAl: Local ancestry and genotype calling uncertainty-aware ancestry-specific allele frequency estimation from admixed samples

**DOI:** 10.1101/2025.07.01.662683

**Authors:** Jiongming Wang, Sebastian Zöllner

## Abstract

In admixed groups, it is of interest to estimate allele frequencies of their ancestral contributing populations. These ancestry-specific allele frequencies inform genetic drivers of disease etiologies, facilitate genome-wide association study interpretations, enhance polygenic risk prediction and portability, and provide insights into demographic history. Their estimation leverages inferred locus-specific ancestry background, i.e., local ancestry, generated by tools like RFMix. However, existing estimation methods lose accuracy and incur biases by failing to model uncertainty from upstream local ancestry inference and genotype calling. Here, we introduce LoGicAl, a novel likelihood-based method for estimating ancestry-specific allele frequencies from admixed samples, simultaneously accommodating uncertainty from ancestry calling, genotyping, and statistical phasing. We demonstrate that modeling these uncertainties substantially reduces estimation errors, resulting in superior accuracy for both sequence-based and array-based genotyping with different levels of local ancestry inference quality. By integrating an accelerated fixed-point algorithm, LoGicAl achieves high scalability and enhanced computational efficiency compared with existing approaches. Applying LoGicAl to admixed cohorts in the 1000 Genomes Project, we illustrate the benefits of local-ancestry-based allele frequency estimates. Together, LoGicAl contributes to genomic analyses of admixed samples by providing precise and rapid ancestry-specific allele frequency estimates, and to constructing the spatial landscape and dynamics of genetic variations in admixed populations at a finer scale.

## 1 Introduction

Accurate allele frequency estimation is crucial to genetic analyses, providing insights into population demographic histories[1], informing population-enriched genetic drivers of disease etiologies[2, 3], facilitating genome-wide association study (GWAS) interpretations and replication cohort designs[4], as well as portability of polygenic risk scores (PRS)[5, 6] in both single-ancestry populations and admixed populations.

Interpretation of allele frequency estimates of admixed populations is complicated by their mosaic structure of genetic ancestral backgrounds[7]. Disregarding this structure when estimating allele frequencies treats genetic variation in current admixed population sample as the mean of the ancestral population allele frequencies weighted by their ancestral contributions. This creates challenges for interpreting admixed populations: since admixed groups can differ by ancestral contributions, allele frequency estimates are not transferable from one group to another. Additionally, it is often of interest to study the pre-admixture ancestral populations. To overcome these challenges, one can estimate the allele frequencies of the ancestral populations from the admixed sample that they contributed to, referred to as ancestry-specific allele frequencies[8].

This approach has been broadly used in genetic analyses of admixed populations. Ancestry-specific allele frequencies have been leveraged to identify ancestry-differentiated risk alleles for complex traits[9], such as asthma[10], blood pressure[11], and mild cognitive impairment[12]. In GWAS design, integrating information from ancestry-specific allele frequencies and matching them across corresponding ancestry cohorts can guide replication designs for validating discovered ancestry-enriched risk variants and improve polygenic prediction accuracy for admixed populations[13]. Moreover, when inferring population structure from admixed samples, incorporating reliable reference ancestry-specific allele frequencies can benefit statistical and computational performance of inference tools[14]. The performance of all these applications depends on the precision of the estimated ancestry-specific allele frequencies.

Several methods have been developed to deconstruct allele frequencies of the ancestral populations for admixed samples. Some methods primarily aim to infer population structure, characterized by admixture proportions (i.e., global ancestry)[15–20]. These methods usually apply mixture models to inform admixture proportion estimates while treating ancestry-specific allele frequencies as nuisance parameters. Another type of methods are explicitly designed to estimate ancestry-specific allele frequencies[21, 22]. These methods integrate information from locus-specific ancestry background (i.e., local ancestry). In heterozygous individuals who are also heterozygous for local ancestry, the ancestry–allele phase is unknown, inducing ambiguity about the ancestral background of each allele. ASAFE[21] estimates allele frequencies by resolving this phase ambiguity through an expectation-maximization (EM) framework; AFA[22] provides allele frequency estimates by modeling the weighted average of ancestry-specific allele frequencies given inferred local (or global) ancestry proportions. These “local-ancestry-aware” methods leverage called (i.e., “best-guess”) local ancestries and genotypes from upstream analyses, ignoring the inherent uncertainty in the ancestry and genotype calling steps. With error-free ancestry calling and genotyping, ASAFE and AFA provide precise estimates. However, both ancestry calling and genotyping can suffer from high uncertainty (or high error rates). For example, continent-level local ancestry estimates have miscall rates of ~5%, rising over 10% in more challenging scenarios[23], and low-coverage sequencing can generate genotype calls with high variance[24]. Directly using such error-prone upstream outputs in existing ancestry-specific allele frequency estimation tools violates their model assumptions and may introduce estimation error and bias.

In this article, we develop a novel ancestry-specific allele frequency estimation method, LoGicAl (Local ancestry and Genotype calling uncertainty-aware ancestry-specific Allele frequency estimation). LoGicAl is an accelerated EM algorithm robust to uncertainty from upstream local ancestry calling and genotype calling. It informs ancestry-specific allele frequency estimates by iteratively calculating sample-wide allele dosages integrated by probabilistic representations of local ancestry and genotype likelihoods over allele frequency spectrum. Variations of LoGicAl are derived to accommodate more flexible input formats. In particular, we provide rapid implementations for phased genotype inputs. We demonstrate the accuracy and efficiency of LoGicAl and benchmark its performance against other state-of-art estimation methods through extensive simulation studies and real data applications.

## 2 Methods

### 2.1 Overview

Here, we present LoGicAl, designed to estimate ancestry-specific allele frequencies from admixed samples by leveraging uncertainty measurement from ancestry calling and genotyping to fine-tune the estimates (Figure 1). LoGicAl is a fully parametric likelihood-based model that uses a hierarchical mixture structure to incorporate the uncertainty from the upstream procedures. We decompose the complete-data likelihood, motivated by this hierarchical mixture model, into conditionally independent components that correspond to different sources of uncertainty. LoGicAl calculates the maximum likelihood estimates using an accelerated EM algorithm. LoGicAl can accommodate either probabilistic representations or best-guess calls of local ancestries and genotypes. Alternative versions of this method further leverage phasing information to generate rapid estimates.

**Figure 1:**
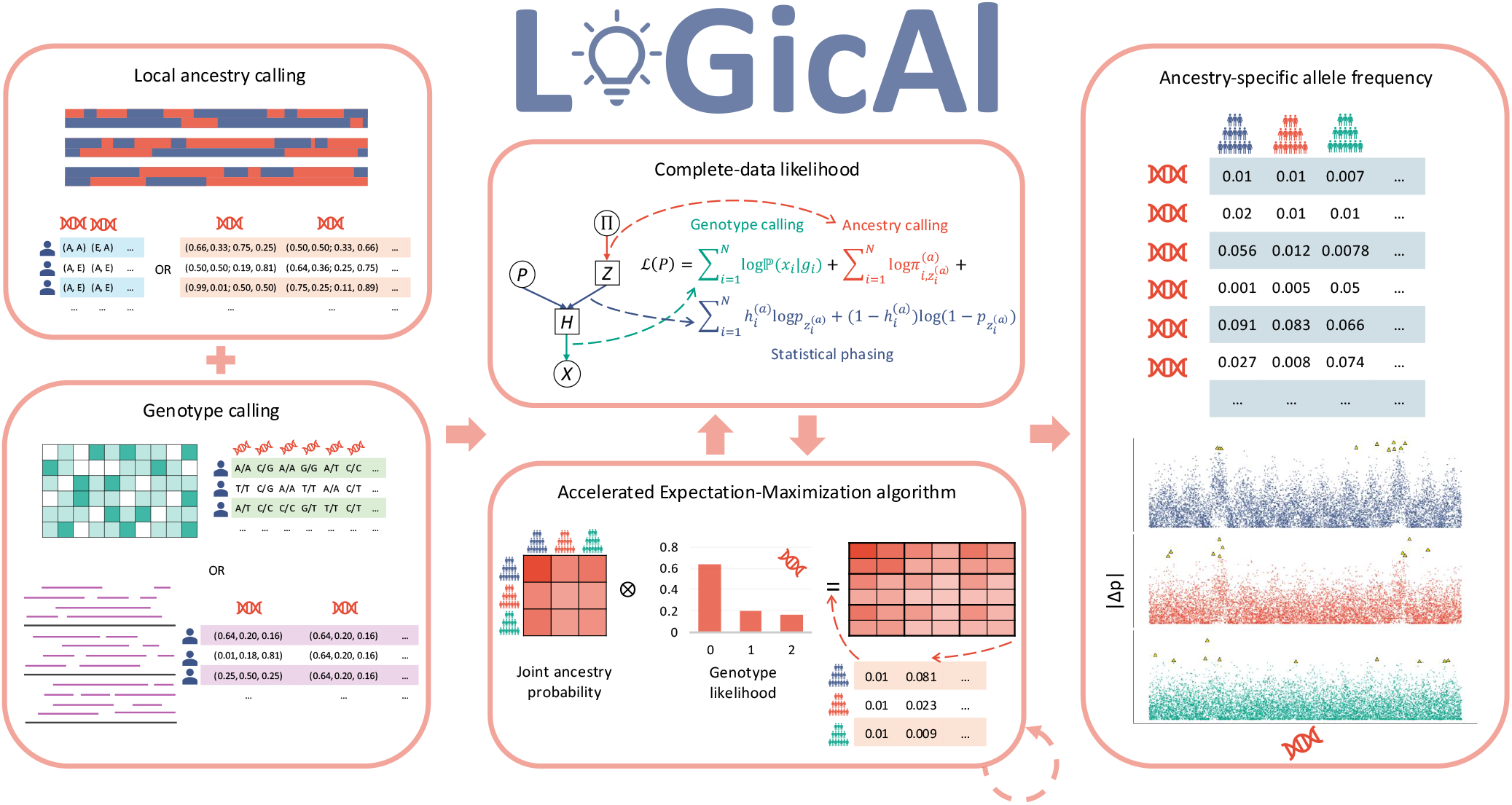
Schematic overview of LoGicAl. LoGicAl estimates ancestry-specific allele frequencies from admixed samples, simultaneously adjusting for uncertainty from upstream local ancestry calling and genotype calling. The inputs of LoGicAl are either best-guess calls or probabilistic representations of local ancestries (upper left) and genotypes (lower left). Specifically, LoGicAl-pl, LoGicAl-al, LoGicAl-pg, and LoGicAl-ag correspond to combinations of local ancestry assignment probabilities (p), or best-guess local ancestries (a), and genotype likelihoods (l), or best-guess genotype calls (g). LoGicAl leverages a hierarchical mixture framework, where the complete-data likelihood is decomposed into conditionally independent parts corresponding to different sources of uncertainty (upper middle). LoGicAl informs parameter estimates using an accelerated EM algorithm: it computes individual-level probabilities for all possible combinations of ancestry-allele phasing and updates allele frequency estimates as sample-wide proportions of ancestry-alternate-allele phases (lower middle). As output, LoGicAl provides ancestry-specific allele frequency estimates of each ancestral population at each genomic position of interest and quantifies pairwise allele frequency differences between ancestral populations across the genome (right). Outlier sites (highlighted in yellow) are defined as the top 0.1% most differentiated variants per pair.

### 2.2 Estimation algorithm

Suppose we have *N* admixed individuals genotyped at *J* bi-allelic sites. Each individual’s genome can originate from *K* ancestral population groups. LoGicAl aims to estimate ancestry-specific allele frequencies **p** = (*p*_1_, …, *p*_*K*_) for these *K* ancestral populations at each site of interest (subscript *j* is omitted for simplicity). Local ancestries are assumed to be estimated using existing inference tools, such as RFMix[25] and FLARE[26]. These local ancestry inference methods provide the most likely ancestry assignments and the corresponding probabilities of assigning each ancestral group. These assignment probabilities can be leveraged to calculate the marginal probabilities of ancestral groups for each individual *i*’s two alleles, denoted by 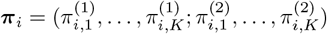. Assuming each individual is unrelated, the observed-data log-likelihood of ancestry-specific allele frequencies is

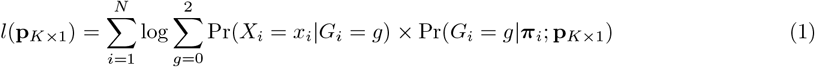

where *X*_*i*_ is observed genotype call (or sequencing reads) and *G*_*i*_ is the true genotype for individual *i*.

LoGicAl maximizes the equation (1) to obtain the maximum likelihood estimates of ancestry-specific allele frequencies. It introduces latent variables 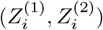 and 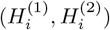, representing the true local ancestry and genotype per chromosome, respectively. Assuming Hardy-Weinberg equilibrium within each ancestral group, LoGicAl estimates ancestry-specific allele frequencies through the following EM algorithm[27] (derivations in Note S1):

#### Algorithm 1

LoGicAl Algorithm for Ancestry-specific Allele Frequency Estimation

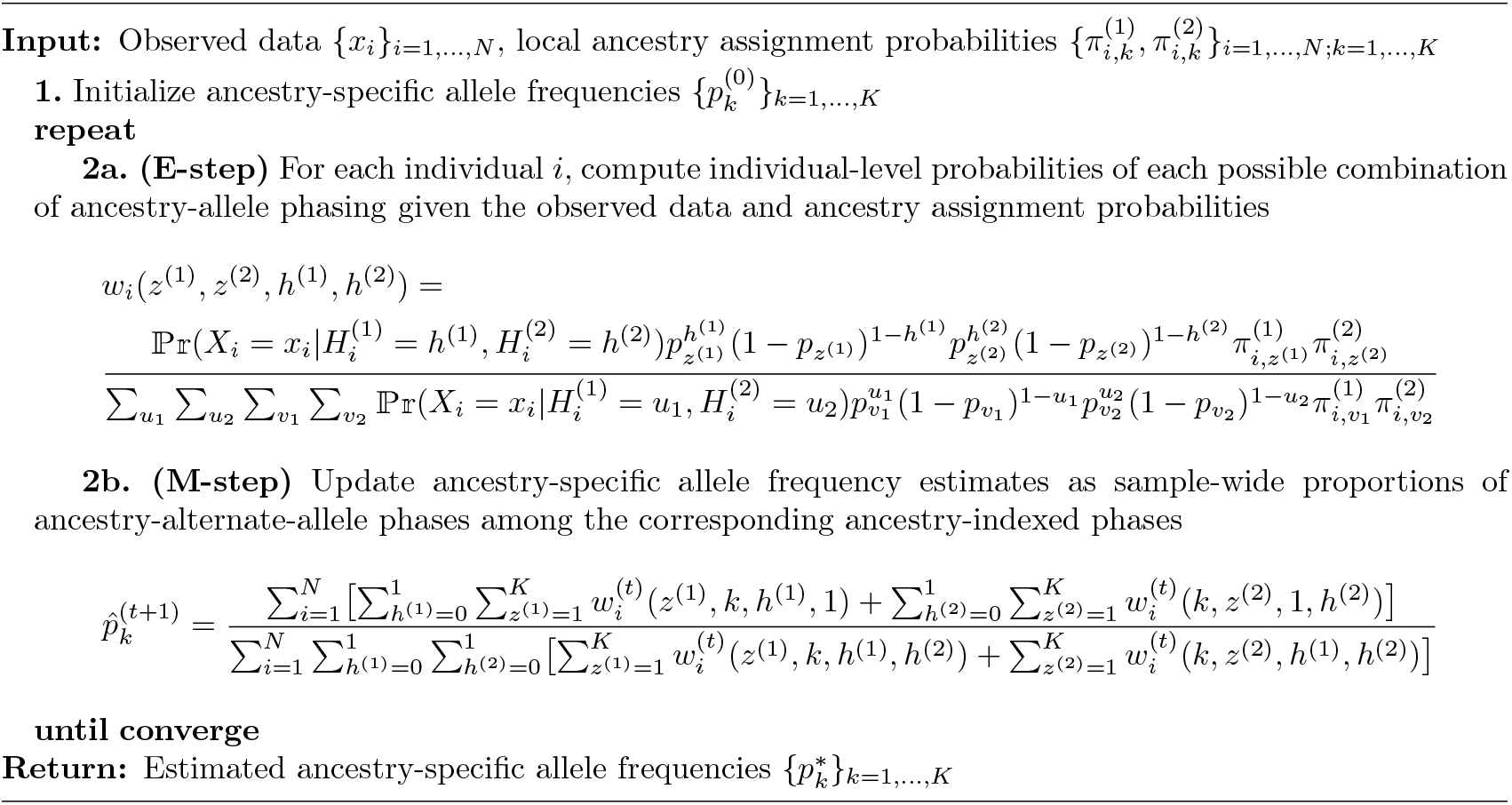

The classical EM algorithm can suffer from slow convergence[28], especially when sample size is large or the uncertainty structure is complicated, thereby limiting its applications to biobank-scale genomic data of admixed samples. We incorporate the squared iterative accelerating technique (SQUAREM)[29, 30], which approximates Newton’s method to tackle the computational burden of the fixed-point algorithms[31], in LoGicAl to enhance its scalability without compromising model preciseness and stability.

### 2.3 Simulation study

We designed a simulation study to investigate estimation accuracy and computation time of LoGicAl and its variations across wide-range settings, and to benchmark with other state-of-art estimation tools (Figure S1). Briefly, we simulated ancestry-specific allele frequencies based on the Balding-Nichols model[32] and specified the probability distribution of underlying local ancestries in each individual. Given the simulated ancestries and ancestry-specific allele frequencies, we sampled the true genotypes and generated observed genotypes from the underlying truth after considering various genotyping error models (details in Note S2). Specifically, we quantified local ancestry resolvability by *α* and fixed *α* = 0.1 across simulation based on its estimates from the 1000 Genomes Project (Table S1). Further, we varied *r* ∈ {0.2, 0.4, 0.6, 0.8, 1} to control ancestral population differentiation: larger *r* approximates allele frequency differentiation observed between populations on different continents (e.g., between San, Han, and Maya populations), whereas smaller *r* reflects differentiation between populations on the same continent[33].

We implemented LoGicAl-pl, LoGicAl-al, LoGicAl-pg, LoGicAl-ag (p: local ancestry assignment probability, a: best-guess local ancestry; l: genotype likelihood, g: best-guess genotype call), ASAFE, and AFA to the simulated unphased genotype inputs. We also applied LoGicAl-pg, LoGicAl-ag, and their phasing-aware variations as well as ASAFE and AFA to the phased genotype inputs. In particular, we implemented ASAFE and AFA using the functions `algorithm_1snp()`and `estimate_frequencies_dynamic_boundary()`with their default parameter settings, except for two modifications in `estimate_frequencies_dynamic_boundary()`: (1) setting `frequency_boundary_grid=c(0.00001,0.01)`, consistent with the AFA Common Workflow Language (CWL) implementation[22], and (2) setting `mac_filter=0`to include rare variants for benchmarking purpose.

### 2.4 Evaluation metrics

We evaluated and summarized the estimation accuracy of different methods using mean squared error (MSE) and relative efficiency (RE). MSE compares the true ancestry-specific allele frequencies **p** = (*p*_1_, …, *p*_*K*_) and their estimates 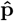 as

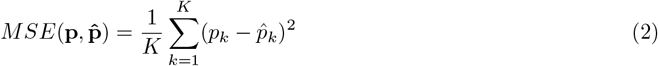

Relative efficiency quantifies the precision of an estimator *T* by comparing the MSE of the empirically-minimal-MSE estimator against that of the estimator *T* as

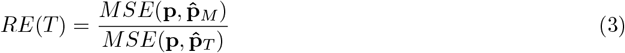

where the minimal-MSE estimates were empirically calculated as point estimates with fully known true local ancestry, genotype, and ancestry-genotype phasing information, thereby reflecting sampling variation only. Therefore, the minimal-MSE estimates correspond to a hypothetical genotyping-error-free sample that has the same sample sizes for each ancestral component as the original sample but consists of independent subsamples, and is thus unaffected by the uncertainty introduced by admixture.

### 2.5 Application to 1000 Genomes Project data

We estimated ancestry-specific allele frequencies from the 1000 Genomes Project (1kGP) cohort[34] using LoGicAl. This dataset contains a total number of 2, 504 unrelated individuals from 26 populations, sequenced at the New York Genome Center. With these deeply-sequenced and phased genotypes, we inferred local ancestries for each individual using RFMix[25] with the option `--node-size=5`. We used the Human Genome Diversity Panel (HGDP) dataset[35, 36] as reference panel for local ancestry calling after pre-processing this reference cohort in alignment with the quality control procedures described in [37]. The 54 populations in HGDP were condensed into seven superpopulations and we used all of them for local ancestry inference: (1) Sub-Saharan Africa, (2) Central & South Asia, (3) East Asia, (4) Europe, (5) Native America, (6) Oceania, and (7) Middle East. After obtaining local ancestry calling results, we applied LoGicAl to calculate ancestry-specific allele frequencies at biallelic single nucleotide polymorphism (SNP) sites from two admixed samples in 1kGP: (1) admixed African sample, including ACB and ASW populations, and (2) admixed American sample, including CLM, MXL, PEL, and PUR populations.

## 3 Results

### 3.1 Estimation of ancestry-specific allele frequencies with uncertainty in ancestry calling and genotyping

To assess the impact of ancestry calling uncertainty and genotyping uncertainty on different estimation methods, we vary the ancestry calling resolution from deterministic, i.e., no uncertainty in local ancestry inference, to probabilistic (*α* = 0.1), and consider genotypes called from hypothetical scenario with no genotyping uncertainty, low-coverage sequencing (1×), and from high-coverage sequencing (30×). We generated the ancestry and genotype information of 100, 000 individuals to approximate the sample size of biobank-scale cohorts, and calculated estimation errors of LoGicAl-pl, LoGicAl-al, LoGicAl-pg, LoGicAl-ag, ASAFE and AFA.

We first examine a scenario with no uncertainty in either ancestry inference or genotype calling (Figure 2, Column 1 and Figures S2-S3). All LoGicAl variations and ASAFE show similar estimates across allele frequency bins. Their MSEs increase monotonously with increasing allele frequency while efficiencies decrease. For the smallest frequency bin, (0.001, 0.005), the MSE is 4.84 × 10^−8^ and the efficiency is 0.915, while for the largest frequency bin, (0.1, 0.5), the MSE is 4.04 × 10^−6^ and the efficiency is 0.769. AFA shows comparable accuracy to LoGicAl and ASAFE for common variants (MAF*>* 0.05), but its accuracy decreases for rare variants.

**Figure 2:**
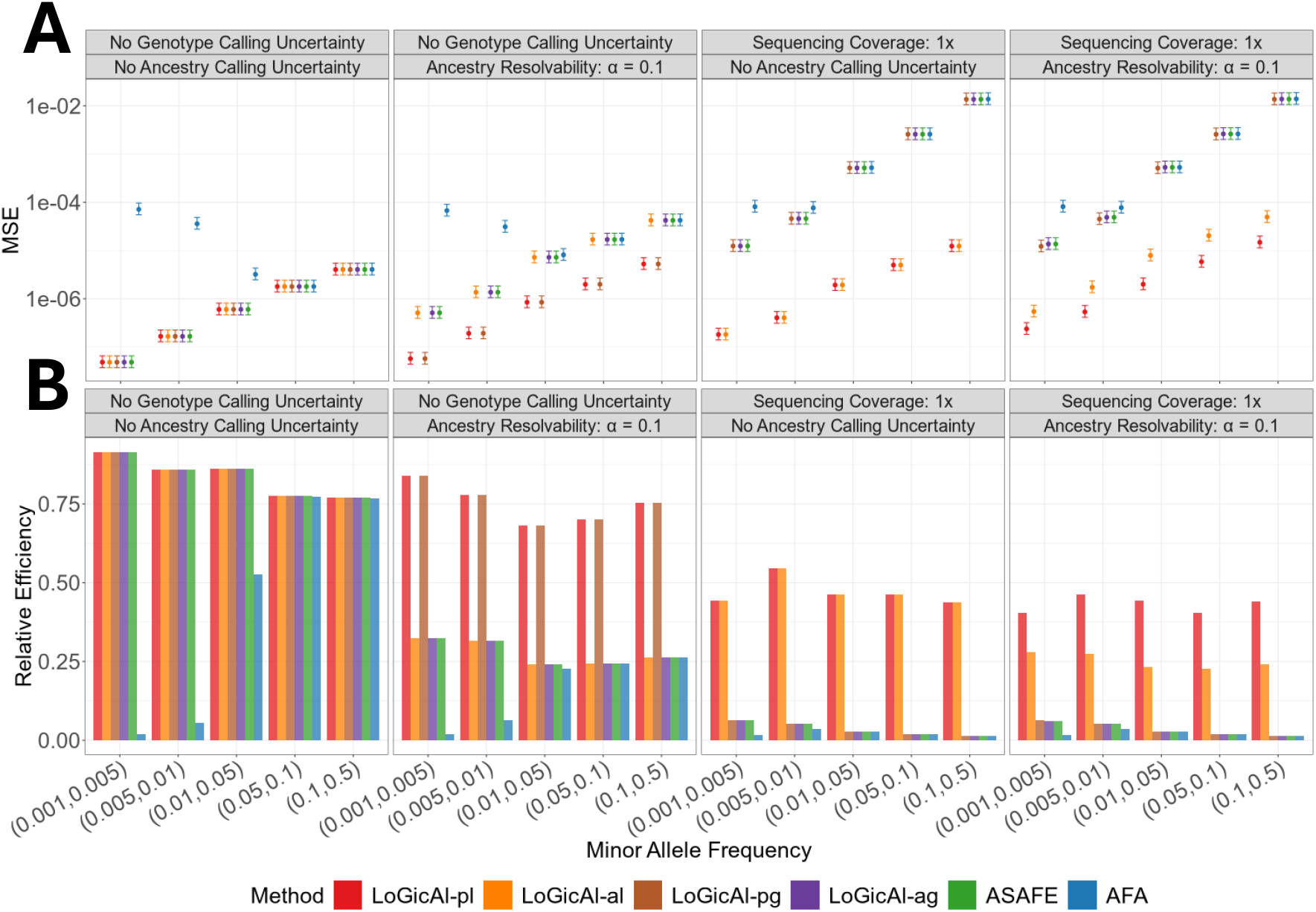
Accuracy of ancestry-specific allele frequency estimation methods across uncertainty scenarios and allele frequency bins. (A) Mean squared error (MSE) and (B) relative efficiency, each calculated over 100 simulation replicates. We simulated 100, 000 unphased individuals from a three-way admixture model with intra-continental ancestral population differentiation (*r* = 0.2). Methods compared include LoGicAl (LoGicAl-pl, LoGicAl-al, LoGicAl-pg, LoGicAl-ag), ASAFE, and AFA under four settings: no calling uncertainty, ancestry calling uncertainty only, genotype calling uncertainty only, and both ancestry and genotype calling uncertainty. In panel A, points denote replicate-averaged MSEs, and intervals indicate 95% confidence intervals for MSEs based on chi-squared approximation. In panel B, bars denote relative efficiencies relative to the optimal estimator.

To evaluate the impact of ancestry calling uncertainty, we consider ancestry inference resolvability *α* = 0.1 while assuming no genotyping uncertainty (Figure 2, Column 2 and Figures S2-S3). Parameter choice of *α* corresponds to the local ancestry posterior distribution estimated from the 1000 Genomes Project (Table S1). Under this scenario, methods that explicitly model ancestry calling uncertainty (LoGicAl-pl/pg) achieve MSEs similar to the scenario without ancestry calling uncertainty, ranging from 5.73 × 10^−8^ for the smallest frequency bin to 5.26 × 10^−6^ for the largest frequency bin. The efficiencies of these methods are slightly lower than under the hypothetical scenario, ranging from 0.841 to 0.680. Methods that do not model ancestry calling uncertainty show ~10-fold higher MSEs: LoGicAl-al/ag and ASAFE range from 5.12 × 10^−7^ to 4.23 × 10^−5^, while AFA ranges from 8.14 × 10^−6^ to 6.73 × 10^−5^. The relative efficiencies of LoGicAl-al/ag and ASAFE drop by 65.6% to 0.324 − 0.242.

To evaluate the impact of genotype calling uncertainty, we consider sequencing depths from low-coverage (1 ×) to high-coverage (30 ×) in the absence of local ancestry inference uncertainty (Figure 2, Column 3 and Figures S2-S3). At 1× depth, methods that explicitly model genotyping uncertainty (LoGicAl-pl/al) achieve MSEs about three times the hypothetical scenario, ranging from 1.82 × 10^−7^ to 1.24 × 10^−5^ across allele frequency bins. The efficiencies of these methods are 43.1% lower, ranging from 0.546 to 0.438. This reduction reflects the uncertainty generated by both low-coverage sequencing and the estimation procedure. Methods that do not model genotyping uncertainty show ~850-fold higher MSEs compared to the scenario without genotyping uncertainty: LoGicAl-pg/ag and ASAFE range from 1.24 ×10^−5^ to 0.0137, while AFA ranges from 7.67 × 10^−5^ to 0.0138. The relative efficiencies of LoGicAl-pg/ag and ASAFE drop by 96.7% to 0.0628 − 0.0138. At 30× depth, estimation accuracy of all methods approaches that under the hypothetical scenario with no uncertainty in either genotyping or ancestry calling (Figures S4-S5).

In a scenario with both uncertain local ancestry calling and low-coverage sequencing (Figure 2, Column 4 and Figures S2-S3), most methods perform similarly to the scenario with genotyping uncertainty only. LoGicAl-pl, which models both types of uncertainty, achieves the lowest MSEs of 2.37 × 10^−7^ 1.48 − 10^−5^ across allele frequency bins. LoGicAl-ag and ASAFE range from 1.37 × 10^−5^ to 0.0138. LoGicAl-pg, modeling ancestry calling uncertainty but not genotyping uncertainty, ranges from 1.22 × 10^−5^ to 0.0137; LoGicAl-al, modeling genotyping uncertainty but not ancestry calling uncertainty, ranges from 5.45 × 10^−7^ to 4.95 × 10^−5^. AFA shows MSEs ranging from 7.77 × 10^−5^ to 0.0139. These performance patterns remain consistent across varying numbers of ancestral populations, from two-way to five-way admixture (Figures S6-S7).

### 3.2 Impact of ancestral population differentiation

We investigate the impact of genetic differentiation among ancestral populations on the precision of ancestry-specific allele frequency estimators in the interplay with uncertainty structure. We varied the differentiation between intra-continental differentiation (*r* = 0.2) and inter-continental differentiation (*r* = 1). Initially focusing on variants with reference allele frequency in (0.01, 0.05), we observe that under the scenario with no ancestry calling or genotyping uncertainty (Figure 3, Column 1), all LoGicAl variations and ASAFE show MSEs ranging from 5.83 × 10^−7^ to 7.05 × 10^−7^, with efficiencies of 0.915 − 0.801. We observe no clear directional effect of increasing ancestral population differentiation. AFA exhibits larger MSE than LoGicAl and ASAFE at *r* = 0.2, and its MSE increases with increasing differentiation. This pattern for AFA is more pronounced for rare variants than for common variants. For common variants with reference allele frequency in (0.1, 0.5), LoGicAl, ASAFE and AFA show similar accuracy across differentiation levels (Figures S2-S3). In the presence of ancestry calling uncertainty (Figure 3, Column 2), MSEs for methods that explicitly model it (LoGicAl-pl/pg) increase slightly, ranging from 6.72 × 10^−7^ to 9.11 × 10^−7^ across differentiation levels, while efficiencies range from 0.776 to 0.680 without clear impact of increasing differentiation. Methods that do not model ancestry calling uncertainty have higher errors that grow with increasing differentiation: LoGicAl-al/ag and ASAFE increase 12-fold at *r* = 0.2 (7.23 × 10^−6^) to 51-fold at *r* = 1 (3.57 × 10^−5^). For these methods efficiencies range from 0.242 at *r* = 0.2 to 0.134 at *r* = 1.

**Figure 3:**
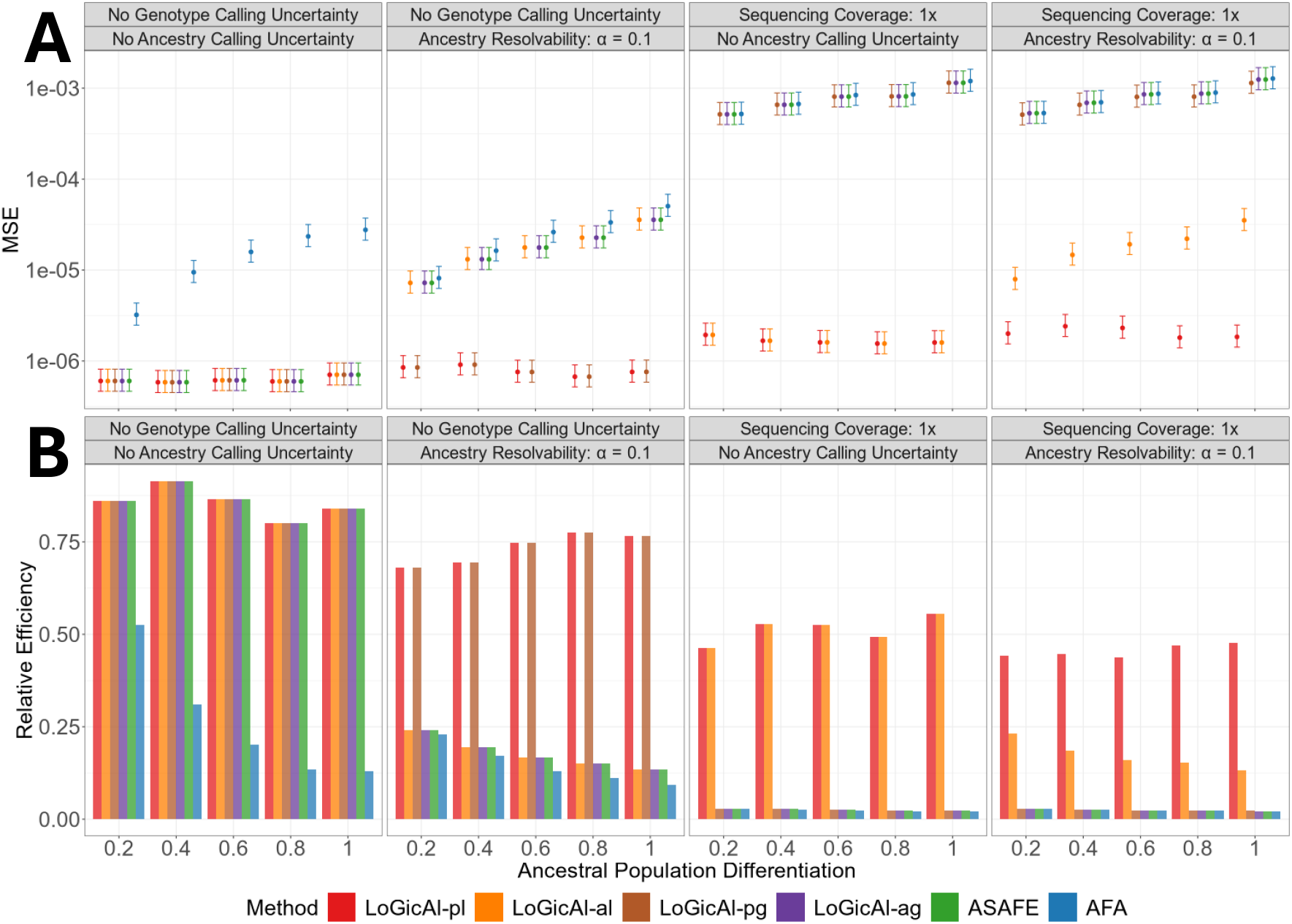
Accuracy of ancestry-specific allele frequency estimation methods across uncertainty scenarios and levels of ancestral population differentiation. (A) Mean squared error (MSE) and (B) relative efficiency, each calculated over 100 simulation replicates. We simulated 100, 000 unphased individuals from a three-way admixture model with reference allele frequency in (0.01, 0.05). Methods compared include LoGicAl (LoGicAl-pl, LoGicAl-al, LoGicAl-pg, LoGicAl-ag), ASAFE, and AFA under four settings: no calling uncertainty, ancestry calling uncertainty only, genotype calling uncertainty only, and both ancestry and genotype calling uncertainty. Smaller value of ancestral population differentiation denotes intra-continental differentiation; larger value denotes inter-continental differentiation. In panel A, points denote replicate-averaged MSEs, and intervals indicate 95% confidence intervals for MSEs based on chi-squared approximation. In panel B, bars denote relative efficiencies relative to the optimal estimator.

In the presence of genotyping uncertainty (Figure 3, Column 3), genotyping-uncertainty-aware methods (LoGicAl-pl/al) achieve MSEs about three times as high as the hypothetical scenario (1.55 × 10^−6^ − 1.93 × 10^−6^), while their efficiencies are reduced to 0.555 − 0.463 without clear impact of increasing differentiation. Genotyping-uncertainty-unaware methods (LoGicAl-pg/ag, ASAFE, AFA) show at least 857 times increased MSEs from 5.16 × 10^−4^ at *r* = 0.2 to 1.15 × 10^−3^ at *r* = 1, with efficiencies below 0.03 for all levels of differentiation. Taken together, MSEs for uncertainty-aware methods remain relatively stable across ancestral population differentiation, while MSEs for uncertainty-unaware methods monotonically increase with greater ancestral differentiation. These performance patterns are consistent across allele frequency bins (Figures S2-S3).

### 3.3 Estimation with phased genotype inputs

To evaluate estimation accuracy when local ancestries and genotypes are jointly phased (Figure 4), we consider a scenario with probabilistic ancestry calling (*α* = 0.1) and a realistic genotyping error rate (*θ* = 5 × 10^−4^). We simulated ancestry and genotype data for 100, 000 individuals and varied the discordance rate from 0% to 10%. This discordance rate quantifies proportions of wrongly flipped ancestry assignments, which can arise from switch errors and double switch errors (i.e., flip errors) in statistical phasing[38] and inferred ancestry segment flips[39] (details in Note S2). We benchmarked LoGicAl-pg and LoGicAl-ag in both phasing-unaware (unphased) and phasing-aware (phased) implementations against ASAFE and AFA.

**Figure 4:**
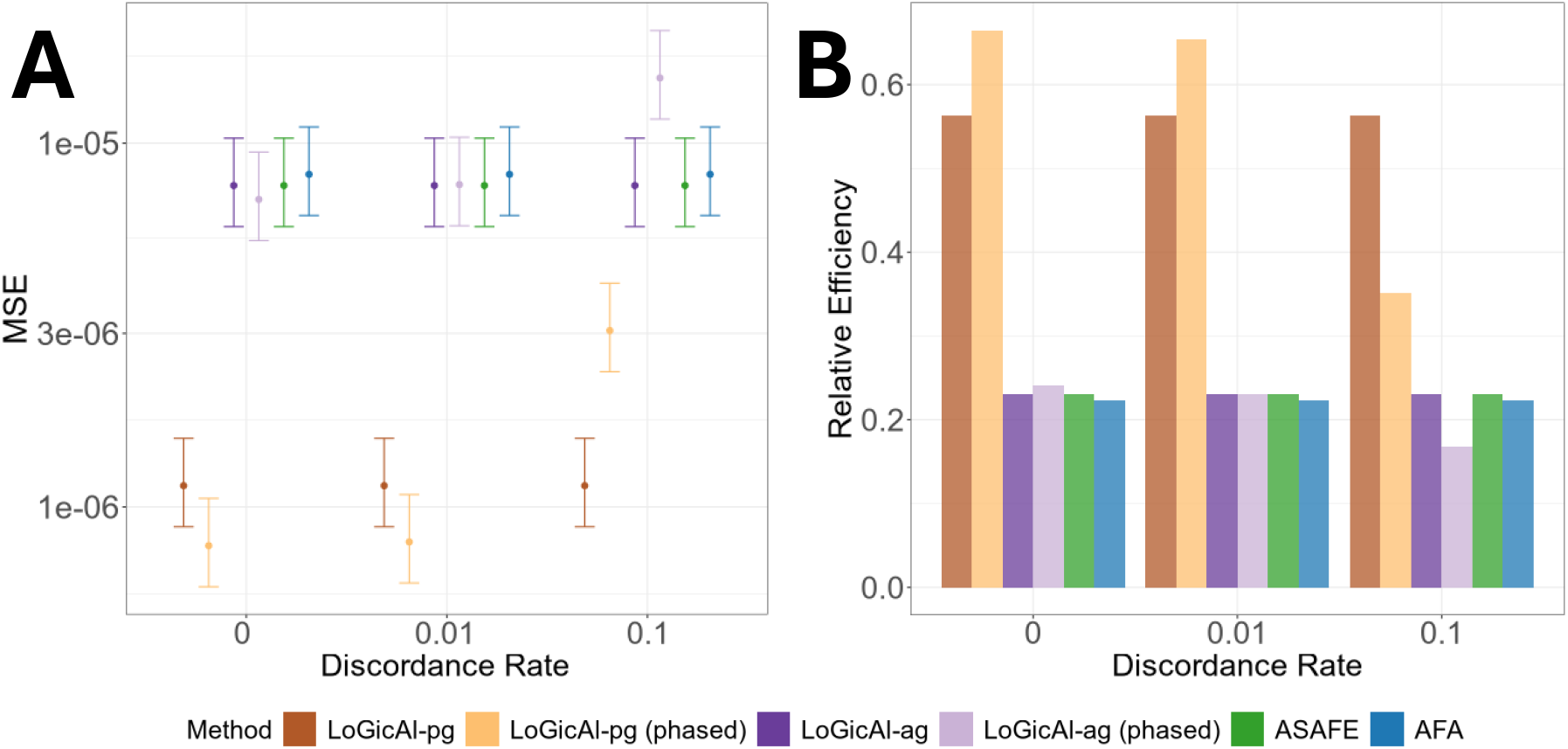
Accuracy of ancestry-specific allele frequency estimation methods across discordance rates. (A) Mean squared error (MSE) and (B) relative efficiency, each calculated over 100 simulation replicates. We simulated 100, 000 phased individuals from a three-way admixture model with reference allele frequency in (0.01, 0.05) and intra-continental differentiation (*r* = 0.2). Methods compared include LoGicAl (LoGicAl-pg, LoGicAl-pg (phased), LoGicAl-ag, LoGicAl-ag (phased)), ASAFE, and AFA under a scenario with probabilistic ancestry calling (*α* = 0.1) and a realistic genotyping error rate (*θ* = 5 × 10^−4^). In panel A, points denote replicate-averaged MSEs, and intervals indicate 95% confidence intervals for MSEs based on chi-squared approximation. In panel B, bars denote relative efficiencies relative to the optimal estimator.

The phasing-unaware methods (LoGicAl-pg/ag, ASAFE, AFA) show no change in MSEs across discordance rates: 1.14 × 10^−6^ for LoGicAl-pg, 7.65 × 10^−6^ for LoGicAl-ag and ASAFE, and 8.21 × 10^−6^ for AFA. In contrast, the phasing-aware methods exhibit increasing errors with higher discordance, resulting in smaller MSEs than the phasing-unaware methods at low discordance ( ~0%) and larger MSEs at high discordance ( ~10%). Specifically, LoGicAl-pg (phased) increases from 7.83 × 10^−7^ at 0% discordance to 3.05 × 10^−6^ at 10% discordance, and LoGicAl-ag (phased) increases from 7.00 × 10^−6^ at 0% discordance to 1.51 × 10^−5^ at 10% discordance. These performance patterns are consistent across allele frequency bins and across levels of ancestral population differentiation (Figures S8-S9), as well as across numbers of ancestral populations (Figures S10-S11).

### 3.4 Computation time

We evaluate the computational performance of LoGicAl, ASAFE, and AFA in terms of runtime for unphased and phased genotype inputs. For unphased data (Table S2), the computation speed of LoGicAl depends on the types of uncertainty that are modeled: modeling both ancestry and genotype calling uncertainty (LoGicAl-pl) requires ~3 − 11 times the computation time of modeling neither (LoGicAl-ag), and modeling either ancestry calling uncertainty (LoGicAl-pg) or genotyping uncertainty (LoGicAl-al) requires ~2 − 3 times and ~1 − 2 times, respectively. Compared to AFA and ASAFE which do not model any uncertainty, LoGicAl is generally substantially faster: compared with AFA, all LoGicAl variants are at least 3 times and up to 56 times faster under the same model. When compared with ASAFE, LoGicAl is up to 6 times faster under the same model or when modeling either ancestry calling uncertainty or genotyping uncertainty. When modeling both, LoGicAl shows similar computation time to ASAFE, ranging from 49.5% faster than ASAFE to 63.4% slower, depending on the complexities of the uncertainty structure. These patterns are consistent across levels of ancestral differentiation (Figure S12).

For phased genotype data (Table S3), the phased version of LoGicAl is ~2 − 5 times faster than the unphased version, and thus uniformly faster than AFA (up to 206 fold) and ASAFE (up to 18 fold). These patterns are consistent across levels of ancestral differentiation (Figure S13).

### 3.5 Application to 1000 Genomes Project

We applied LoGicAl to estimate ancestry-specific allele frequencies from the admixed samples in the 1000 Genomes Project. Specifically, we estimated (1) African and European ancestral allele frequencies in the admixed African sample (ACB and ASW cohorts), and (2) African, European and Native American ancestral allele frequencies in the admixed American sample (CLM, MXL, PEL and PUR cohorts).

To validate LoGicAl estimates, we compared the ancestry-specific allele frequency estimates with allele frequencies of the corresponding non-admixed ancestral populations in HGDP. Correlations are high for ancestry-matched comparisons and lower for ancestry-mismatched pairs: from 1kGP admixed African cohorts, African ancestral estimates highly correlate with HGDP African cohorts (*r* = 0.982), and European ancestral estimates highly correlate with HGDP Europeans (*r* = 0.975). In contrast, cross-ancestry correlations are lower (African ancestral estimates versus HGDP Europeans *r* = 0.792; European ancestral estimates versus HGDP Africans *r* = 0.772). Similarly, from 1kGP admixed American cohorts, correlations of African ancestral allele frequency estimates with HGDP African, European and Native American cohorts are 0.965, 0.760, and 0.702, respectively; correlations of European ancestral estimates with HGDP African, European and Native American cohorts are 0.772, 0.994, and 0.848; correlations of Native American ancestral estimates with HGDP African, European and Native American cohorts are 0.705, 0.837, and 0.985.

To compare the impact of using local versus global ancestry on allele frequency estimation, we calculated correlations between LoGicAl estimates (local-ancestry-based) and allele frequency estimates derived from methods that assign a single global ancestry label per individual, as in the gnomAD v4.1.0 whole-genome sequencing dataset[40]. In gnomAD, both non-admixed and admixed African cohorts are labeled as “African”, and admixed American cohorts are labeled as “American”. Correlations between gnomAD African allele frequencies and LoGicAl African ancestral estimates are 0.989 for 1kGP admixed African cohorts and 0.970 for 1kGP admixed American cohorts; correlation between gnomAD American allele frequencies and LoGicAl Native American ancestral estimates is 0.923.

To illustrate how local-versus global-ancestry-based approaches differ at specific loci, we compared their estimates at LoGicAl-identified ancestry-differentiated variants on chromosome 1 (Figure S14). LoGicAl (local-ancestry-based) provides more distinct estimates of ancestral population differentiation compared to gnomAD (global-ancestry-based). For example, in 1kGP admixed African cohorts, *rs*2814778 (in *ACKR*1) is the most differentiated site with African ancestral estimate of 0.996 and European ancestral estimate of *<* 0.001, whereas gnomAD estimates are AFR = 0.822 and EUR = 0.004. Similar patterns are observed in estimates from 1kGP admixed American cohorts (Table S4).

## 4 Discussion

In this article, we develop LoGicAl, a novel accurate, flexible, and scalable tool for estimating ancestry-specific allele frequencies from admixed samples. LoGicAl applies likelihood-based estimation that resolves an ancestry-genotype-integrated hierarchical mixture structure through an accelerated EM algorithm. LoGicAl provides faster implementations for phased genotype inputs when ancestry-genotype phasing is reliable. By effectively modeling uncertainty arising from ancestry calling and genotyping, as well as accounting for statistical phasing errors, LoGicAl achieves superior accuracy and scalability compared with state-of-art estimation methods in both realistic-parameter-driven simulations and analyses of 1000 Genomes Project data.

LoGicAl accommodates four important features of modern genetic analyses of admixed samples. First, it improves ancestry-specific allele frequency estimation accuracy by explicitly modeling uncertainty in local ancestry inference. Treating local ancestry as probability distributions rather than as point calls is particularly important when inferring local ancestry is challenging, for example, when ancestral populations are closely related[26]. Ignoring such uncertainty induces bias by shrinking ancestry-specific allele frequency estimates towards the unstratified allele frequency. The magnitude of this error grows with increasing ancestral population differentiation. Though local ancestry misspecifications are less common for more differentiated ancestral populations, even rare misspecifications can produce substantial errors if not appropriately modeled.

Second, LoGicAl provides accurate ancestry-specific allele frequency estimates from low-coverage whole-genome sequencing. Low-coverage sequencing has garnered broad interests for its ability to capture novel variants in underrepresented populations[24], increase GWAS power[41] and to improve PRS transferability[42]. Companies like Regeneron use low-pass genome plus high-coverage exome sequencing at scale as a cost-effective alternative to WGS or WES[43]. Our results demonstrate that ignoring genotyping uncertainty inherent to low-coverage sequencing induces bias in ancestry-specific allele frequency estimates. By explicitly modeling this uncertainty, LoGicAl substantially mitigates estimation errors and provides more precise and efficient estimates compared to existing methods across allele frequencies.

Third, phasing-aware LoGicAl reduces ancestry-specific allele frequency estimation errors by leveraging ancestry–genotype phase when phasing error is negligible. With increasing phasing error, phasing-aware estimators become less precise, exceeding MSEs of phasing-unaware estimators at reported phasing error rates in the literature[44]. Practically, the phased implementation is preferred when high-quality ancestry-genotype phasing is available (e.g., trio-based inference), whereas phasing-unaware approaches are preferred when phasing quality is uncertain or unknown.

Fourth, LoGicAl accelerates the implementation of the classical EM algorithm through the squared iterative method, without compromising accuracy or stability. When phased genotype inputs are available, they substantially reduce the parameter space within which the accelerated EM algorithm searches, thereby further reducing computational costs. Altogether, these computational improvements enable the utilities of LoGicAl for analyzing biobank-scale genomic data of admixed samples, such as the TOPMed program[45]. Our application to 1000 Genomes Project admixed samples shows that LoGicAl produces ancestry-specific allele frequency estimates that are well calibrated to non-admixed HGDP references, recovering allele frequencies in the corresponding ancestral populations. Comparing local- and global-ancestry strategies, we find that global labels reflect general ancestry patterns but obscure admixture-specific structure, resulting in shrunken allele frequency estimates. In contrast, the local-ancestry approach pinpoints locus-specific ancestral population differentiation masked with global labeling. We illustrate this by, for example, *rs*2814778 in the *ACKR*1 region, which codes atypical chemokine receptor 1 and can affect malaria susceptibility[46]. At this variant, African and European ancestral population differentiation is more pronounced using LoGicAl-derived estimates than with global-ancestry-based frequencies from gnomAD.

Several extensions could further enhance statistical and computational performance of LoGicAl. First, the current estimation framework assumes unrelated samples. While point estimators remain unbiased under relatedness, dependence can inflate precision metrics if not accounted for. Robust estimators or kinship-aware weighting could be leveraged to inform calibrated standard errors and confidence intervals. Second, LoGicAl provides marginal site-wise ancestry-specific allele frequency estimates and it does not explicitly leverage linkage disequilibrium (LD) structure across sites. Grouping variants by LD structure and local ancestry segments and performing block-wise analyses could benefit its computational performance[47].

To summarize, LoGicAl provides comprehensive and precise descriptions of genetic variation across genome and informs ancestry-specific allele frequency spectrum for admixed populations. LoGicAl further contributes to constructing a precise spatial landscape and dynamics of genetic variation across geographical regions, and promote the understandings in population structure and demographic history at a finer scale.

## Data and code availability

The LoGicAl software is available at https://github.com/student-jw/LoGicAl.

## Acknowledgments

This work was supported by the National Institutes of Health (NIH) grants R01 HG011031 and R01 HG005855. We acknowledge Gonçalo Abecasis for helpful discussion on phasing-aware approaches.

## Supplemental figures

**Figure S1:**
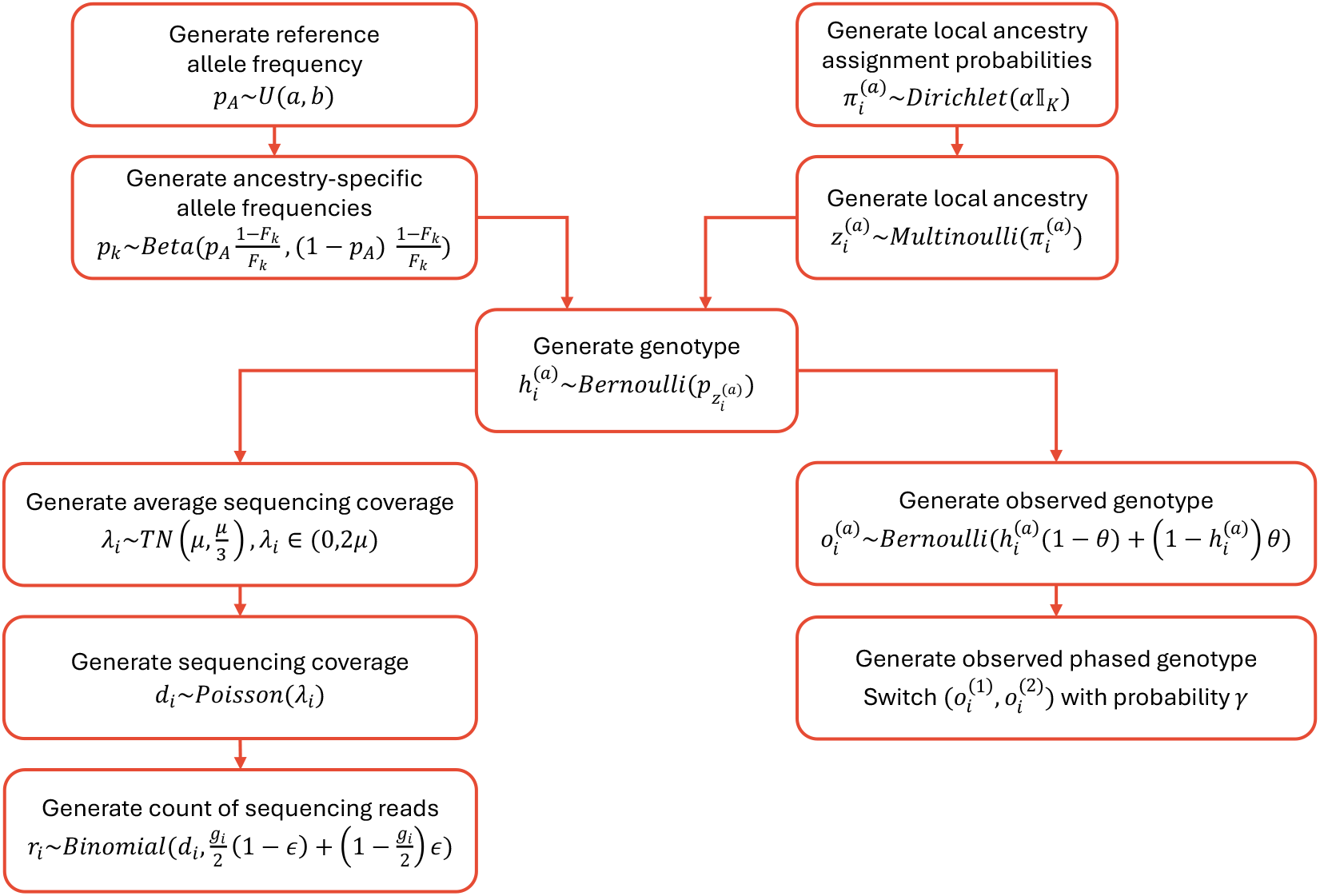
Simulation overview.

**Figure S2:**
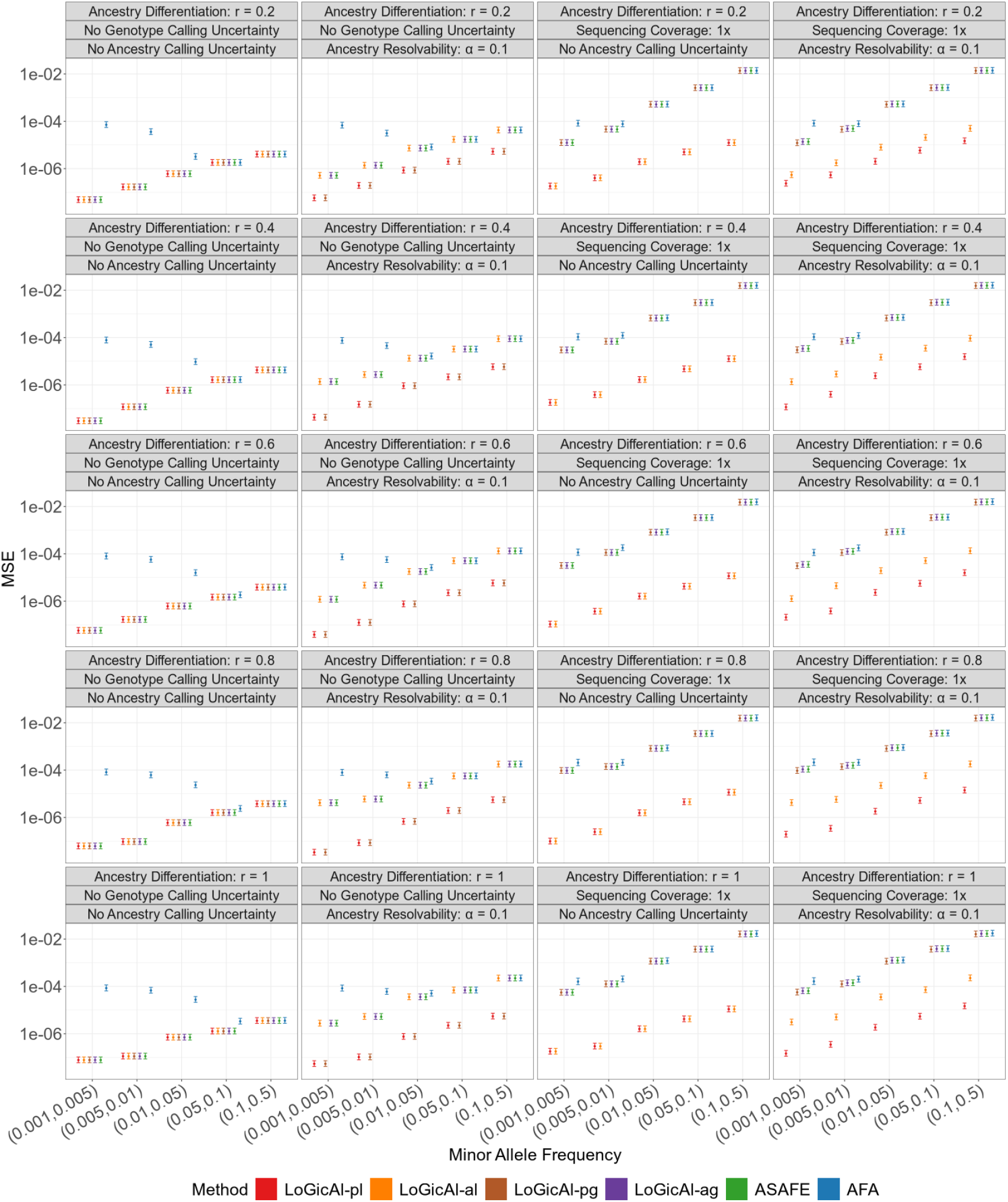
Mean squared error of ancestry-specific allele frequency estimation methods across uncertainty scenarios, allele frequency bins and levels of ancestral population differentiation. We simulated 100, 000 unphased individuals from a three-way admixture model with intra-continental differentiation (*r* = 0.2) to inter-continental differentiation (*r* = 1). Methods compared include LoGicAl (LoGicAl-pl, LoGicAl-al, LoGicAl-pg, LoGicAl-ag), ASAFE, and AFA under four settings: no calling uncertainty, ancestry calling uncertainty only, genotype calling uncertainty only, and both ancestry and genotype calling uncertainty. Points denote replicate-averaged MSEs, and intervals indicate 95% confidence intervals for MSEs based on chi-squared approximation.

**Figure S3:**
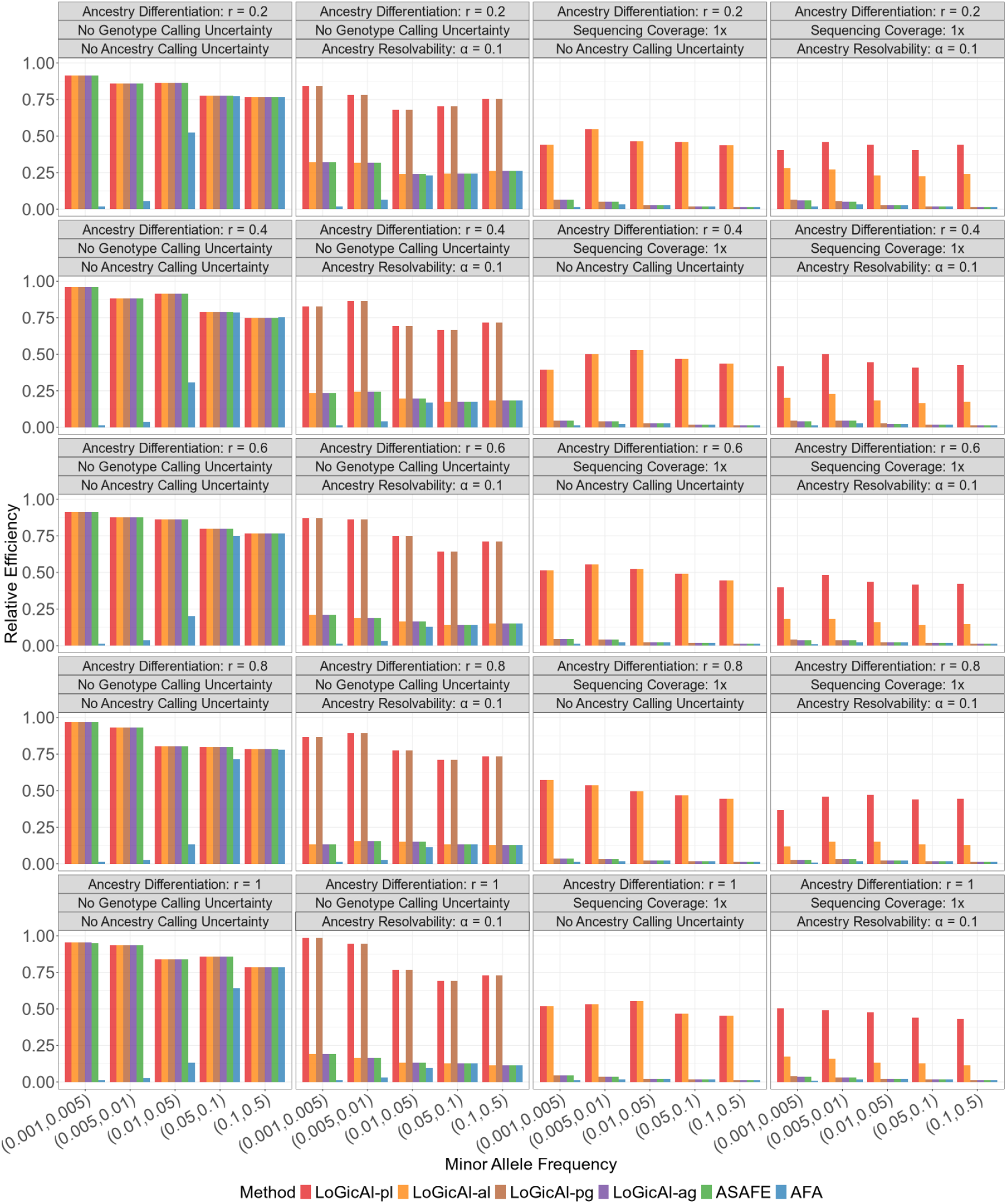
Relative efficiency of ancestry-specific allele frequency estimation methods across uncertainty scenarios, allele frequency bins and levels of ancestral population differentiation. We simulated 100, 000 unphased individuals from a three-way admixture model with intra-continental differentiation (*r* = 0.2) to inter-continental differentiation (*r* = 1). Methods compared include LoGicAl (LoGicAl-pl, LoGicAl-al, LoGicAl-pg, LoGicAl-ag), ASAFE, and AFA under four settings: no calling uncertainty, ancestry calling uncertainty only, genotype calling uncertainty only, and both ancestry and genotype calling uncertainty. Bars denote relative efficiencies relative to the optimal estimator.

**Figure S4:**
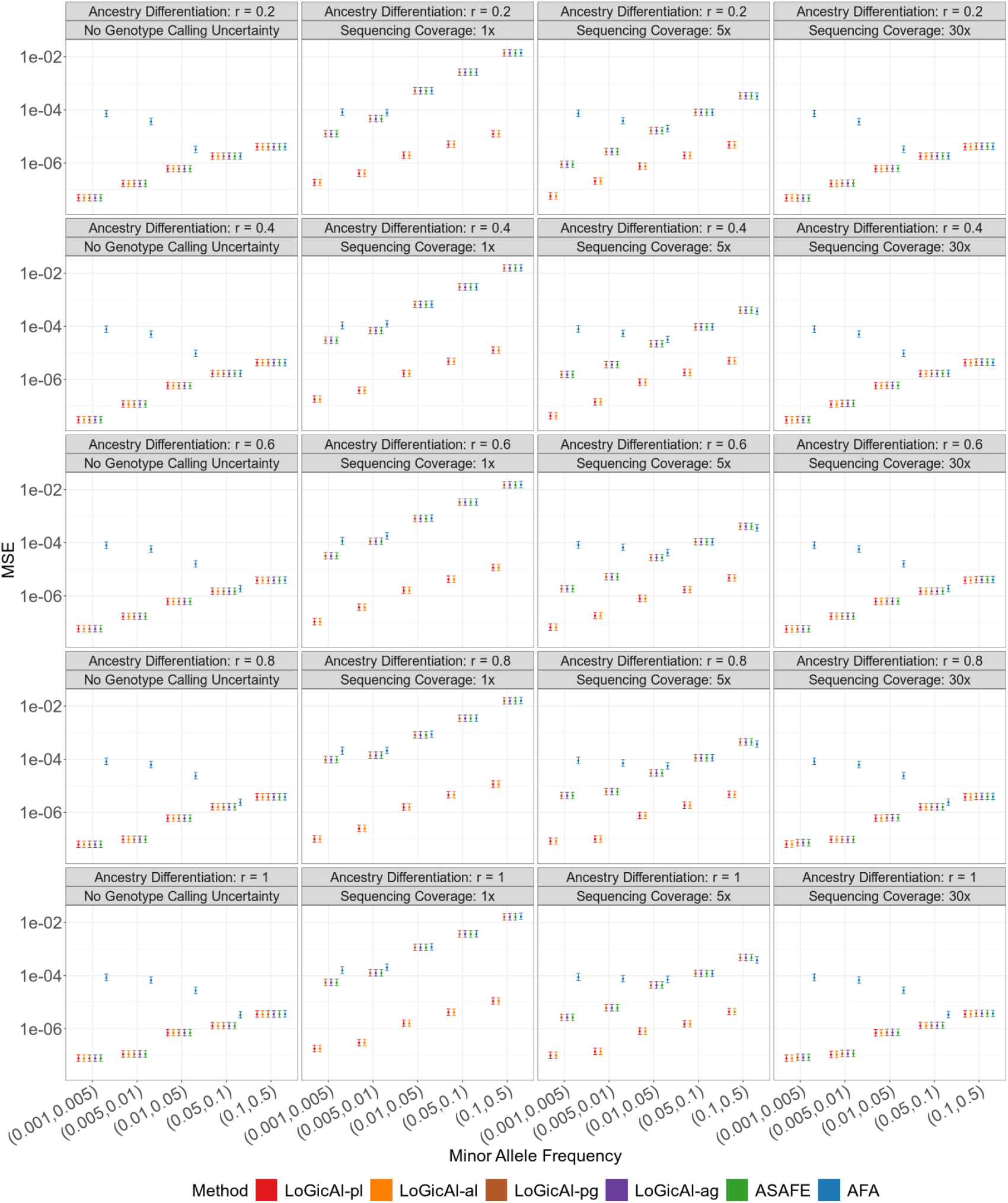
Mean squared error of ancestry-specific allele frequency estimation methods across sequencing depths, allele frequency bins and levels of ancestral population differentiation. We simulated 100, 000 unphased individuals from a three-way admixture model with intra-continental differentiation (*r* = 0.2) to inter-continental differentiation (*r* = 1) while assuming no ancestry inference uncertainty. Methods compared include LoGicAl (LoGicAl-pl, LoGicAl-al, LoGicAl-pg, LoGicAl-ag), ASAFE, and AFA. Points denote replicate-averaged MSEs, and intervals indicate 95% confidence intervals for MSEs based on chi-squared approximation.

**Figure S5:**
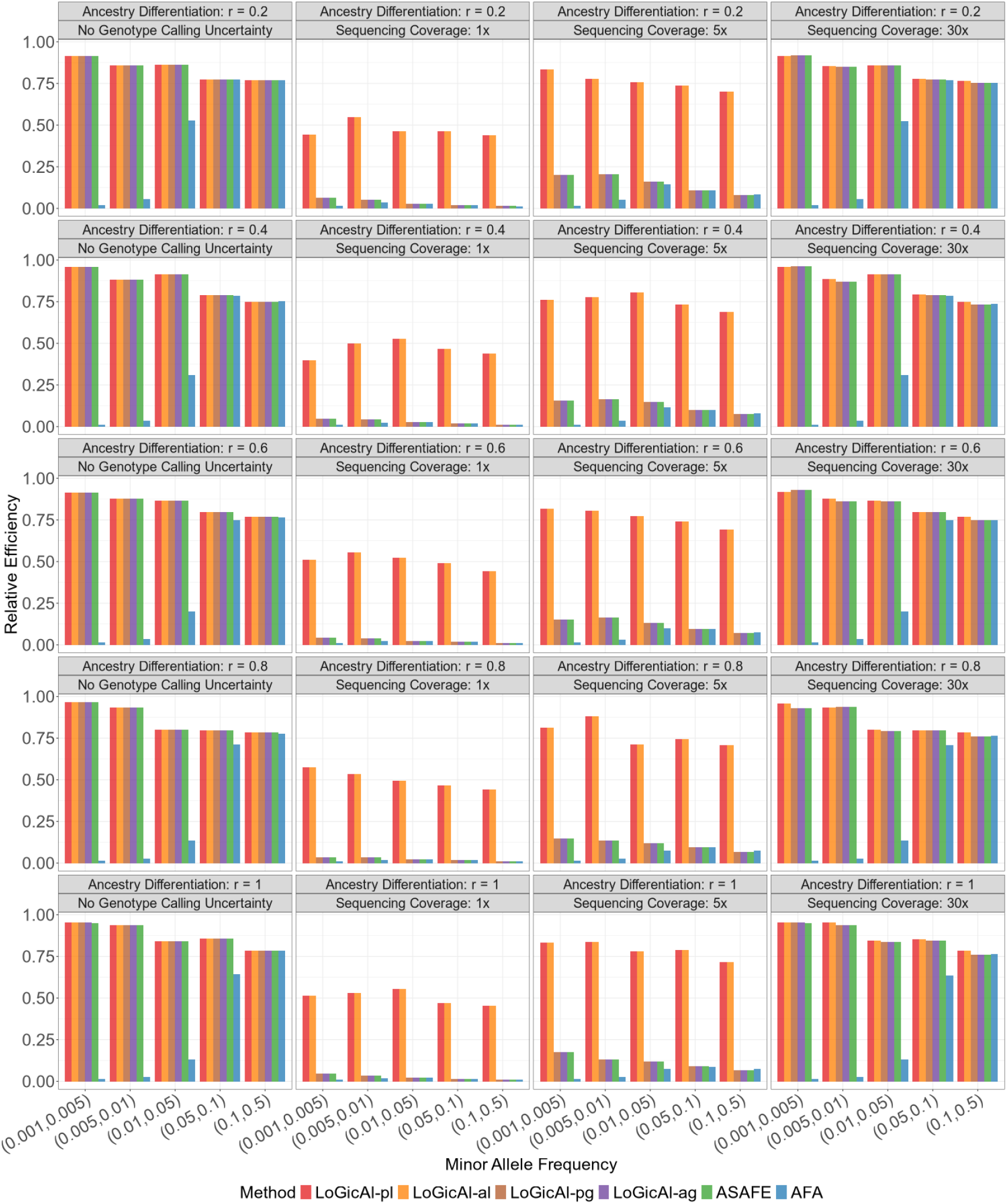
Relative efficiency of ancestry-specific allele frequency estimation methods across sequencing depths, allele frequency bins and ancestral population differentiation. We simulated 100, 000 unphased individuals from a three-way admixture model with intra-continental differentiation (*r* = 0.2) to inter-continental differentiation (*r* = 1) while assuming no ancestry inference uncertainty. Methods compared include LoGicAl (LoGicAl-pl, LoGicAl-al, LoGicAl-pg, LoGicAl-ag), ASAFE, and AFA. Bars denote relative efficiencies relative to the optimal estimator.

**Figure S6:**
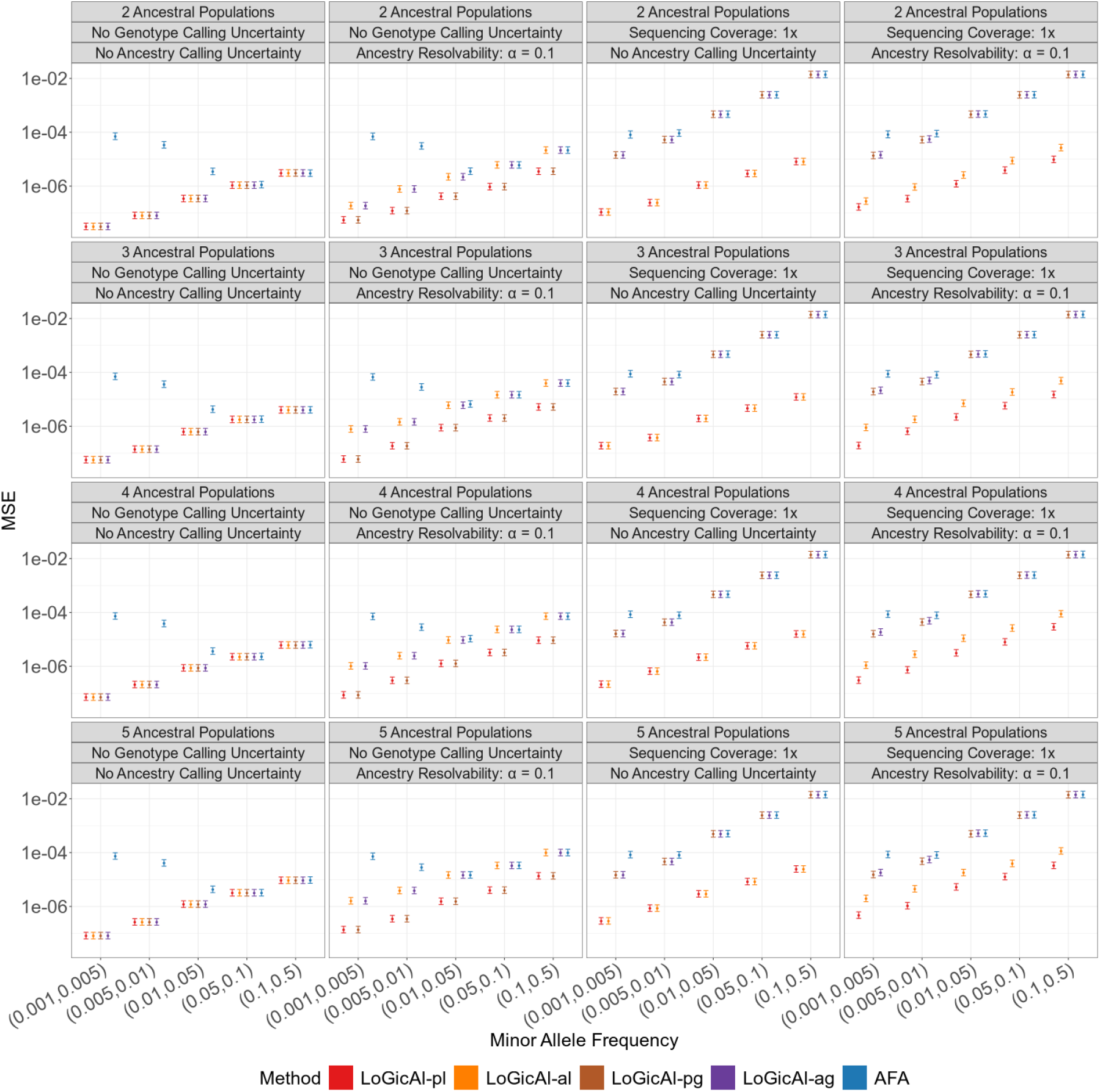
Mean squared error of ancestry-specific allele frequency estimation methods across uncertainty scenarios, allele frequency bins and numbers of ancestral populations. We simulated 100, 000 unphased individuals under two-to five-way admixture with intra-continental differentiation. Methods compared include LoGicAl (LoGicAl-pl, LoGicAl-al, LoGicAl-pg, LoGicAl-ag) and AFA under four settings: no calling uncertainty, ancestry calling uncertainty only, genotype calling uncertainty only, and both ancestry and genotype calling uncertainty. ASAFE is not included as it is developed for three-way (and two-way) admixture scenario only. Points denote replicate-averaged MSEs, and intervals indicate 95% confidence intervals for MSEs based on chi-squared approximation.

**Figure S7:**
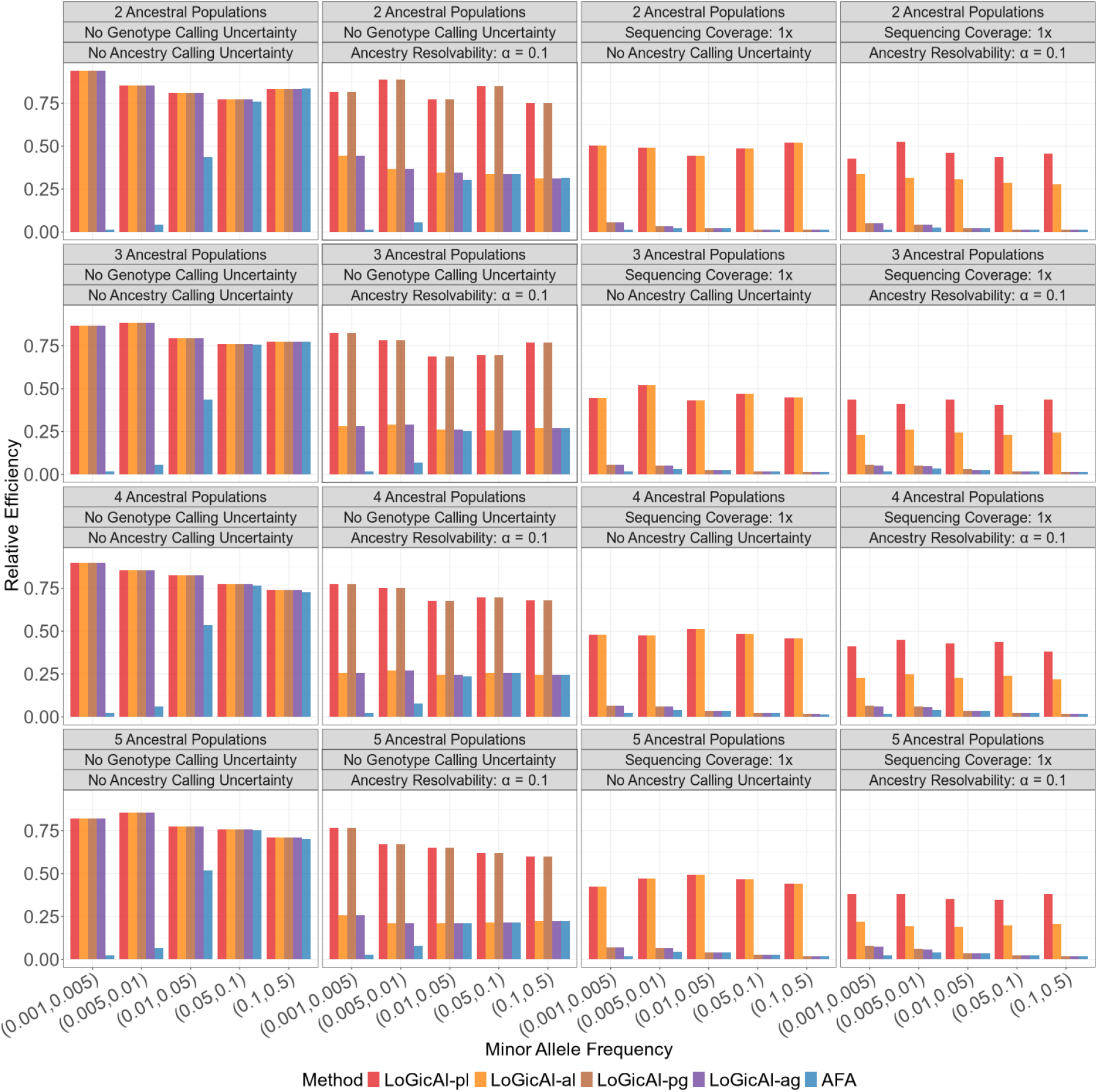
Relative efficiency of ancestry-specific allele frequency estimation methods across uncertainty scenarios, allele frequency bins and numbers of ancestral populations. We simulated 100, 000 unphased individuals under two-to five-way admixture with intra-continental differentiation. Methods compared include LoGicAl (LoGicAl-pl, LoGicAl-al, LoGicAl-pg, LoGicAl-ag) and AFA under four settings: no calling uncertainty, ancestry calling uncertainty only, genotype calling uncertainty only, and both ancestry and genotype calling uncertainty. ASAFE is not included as it is developed for three-way (and two-way) admixture scenario only. Bars denote relative efficiencies relative to the optimal estimator.

**Figure S8:**
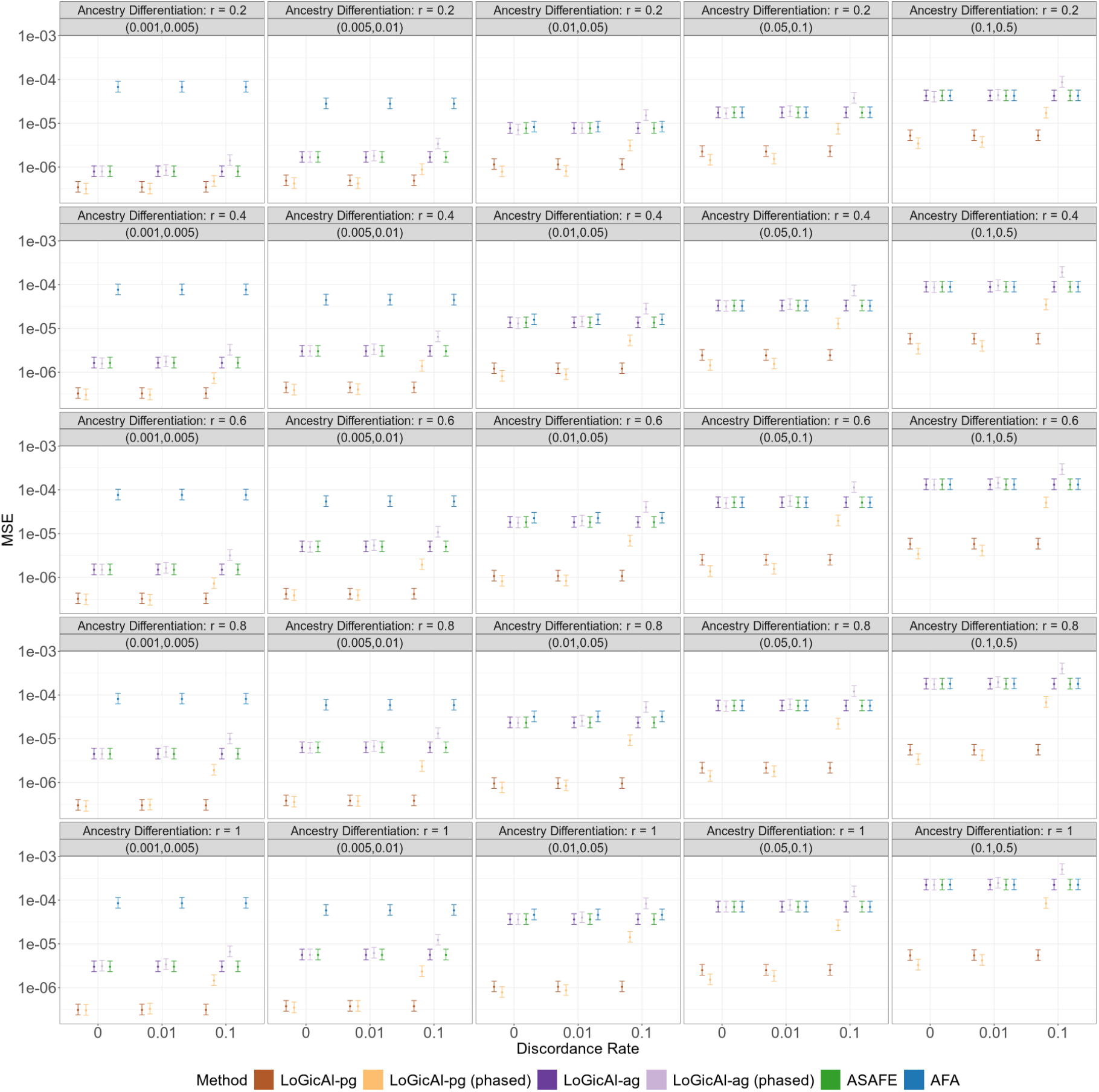
Mean squared error of ancestry-specific allele frequency estimation methods across discordance rates, allele frequency bins and levels of ancestral population differentiation. We simulated 100, 000 phased individuals from a three-way admixture model under a scenario with probabilistic ancestry calling (*α* = 0.1) and a realistic genotyping error rate (*θ* = 5 × 10^−4^). Methods compared include LoGicAl (LoGicAl-pg, LoGicAl-pg (phased), LoGicAl-ag, LoGicAl-ag (phased)), ASAFE and AFA. Points denote replicate-averaged MSEs, and intervals indicate 95% confidence intervals for MSEs based on chi-squared approximation.

**Figure S9:**
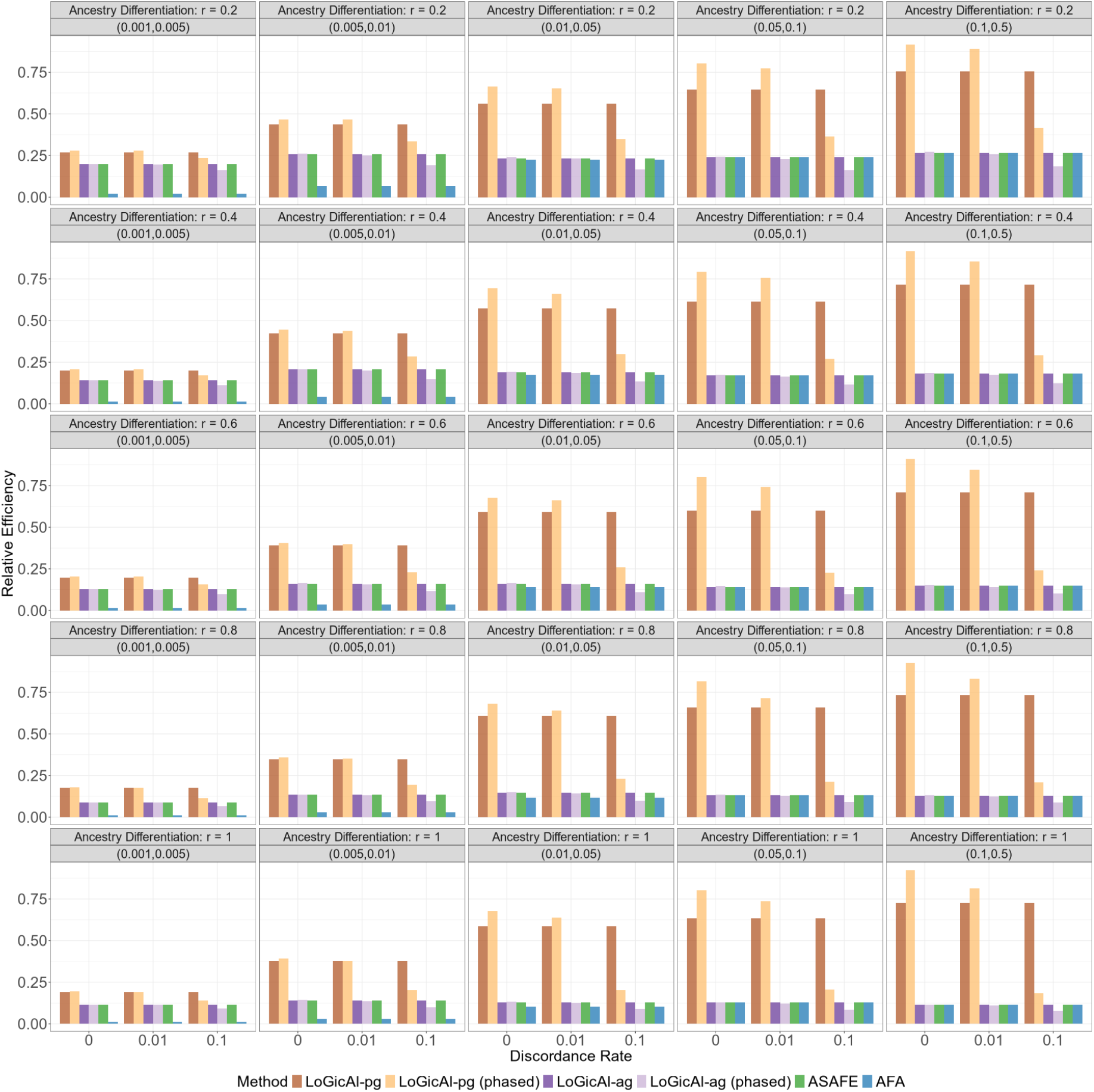
Relative efficiency of ancestry-specific allele frequency estimation methods across discordance rates, allele frequency bins and levels of ancestral population differentiation. We simulated 100, 000 phased individuals from a three-way admixture model under a scenario with probabilistic ancestry calling (*α* = 0.1) and a realistic genotyping error rate (*θ* = 5 × 10^−4^). Methods compared include LoGicAl (LoGicAl-pg, LoGicAl-pg (phased), LoGicAl-ag, LoGicAl-ag (phased)), ASAFE and AFA. Bars denote relative efficiencies relative to the optimal estimator.

**Figure S10:**
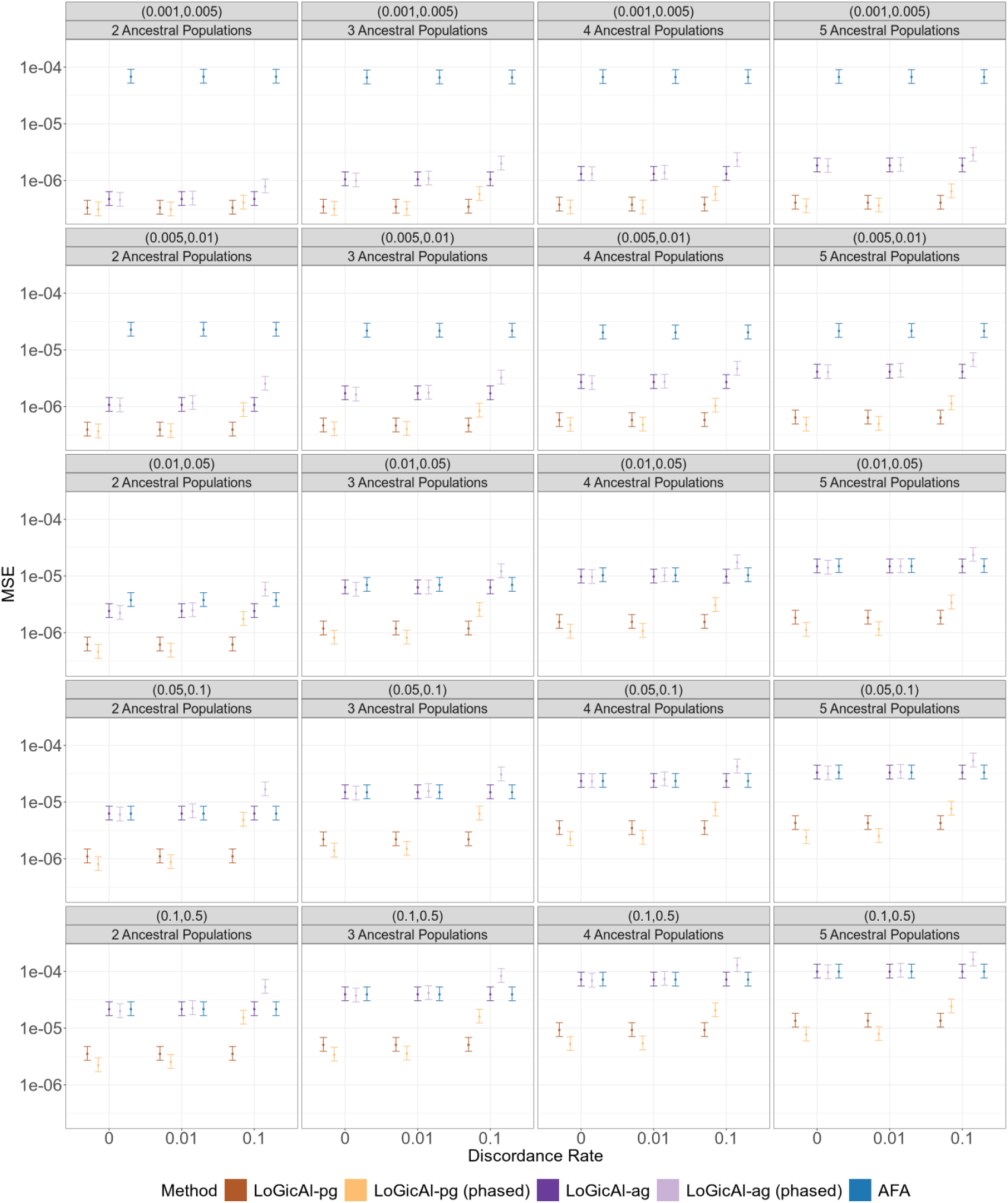
Mean squared error of ancestry-specific allele frequency estimation methods across discordance rates, allele frequency bins and numbers of ancestral populations. We simulated 100, 000 phased individuals under two-to five-way admixture with intra-continental differentiation. Methods compared include LoGicAl (LoGicAl-pg, LoGicAl-pg (phased), LoGicAl-ag, LoGicAl-ag (phased)) and AFA under a scenario with probabilistic ancestry calling (*α* = 0.1) and a realistic genotyping error rate (*θ* = 5 × 10^−4^). ASAFE is not included as it is developed for three-way (and two-way) admixture scenario only. Points denote replicate-averaged MSEs, and intervals indicate 95% confidence intervals for MSEs based on chi-squared approximation.

**Figure S11:**
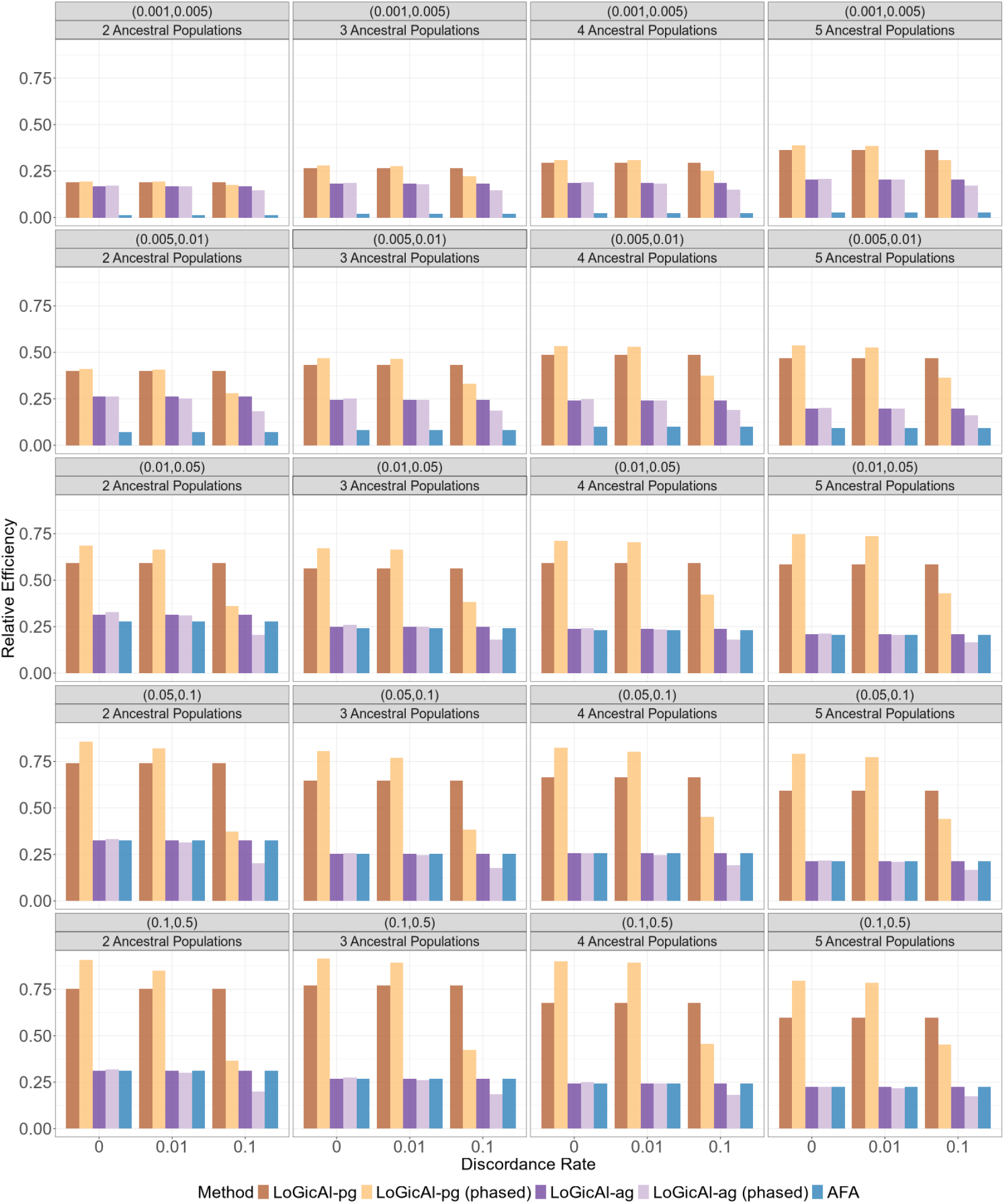
Relative efficiency of ancestry-specific allele frequency estimation methods across discordance rates, allele frequency bins and numbers of ancestral populations. We simulated 100, 000 phased individuals under two-to five-way admixture with intra-continental differentiation. Methods compared include LoGicAl (LoGicAl-pg, LoGicAl-pg (phased), LoGicAl-ag, LoGicAl-ag (phased)) and AFA under a scenario with probabilistic ancestry calling (*α* = 0.1) and a realistic genotyping error rate (*θ* = 5 × 10^−4^). ASAFE is not included as it is developed for three-way (and two-way) admixture scenario only. Bars denote relative efficiencies relative to the optimal estimator.

**Figure S12:**
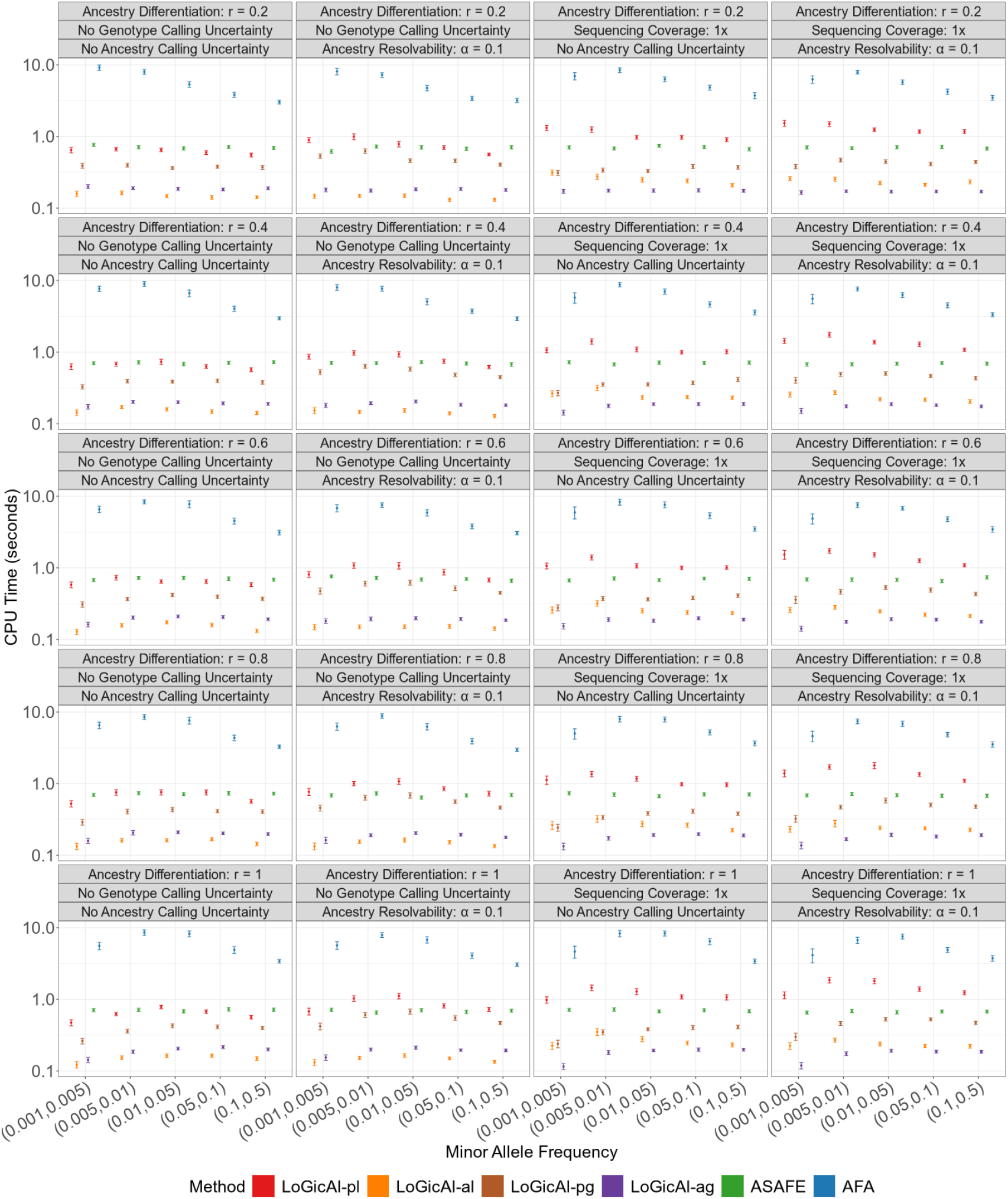
CPU runtime (seconds) of ancestry-specific allele frequency estimation methods across uncertainty scenarios, allele frequency bins and levels of ancestral population differentiation. We simulated 100, 000 unphased individuals from a three-way admixture model with intra-continental differentiation (*r* = 0.2) to inter-continental differentiation (*r* = 1). Methods compared include LoGicAl (LoGicAl-pl, LoGicAl-al, LoGicAl-pg, LoGicAl-ag), ASAFE, and AFA under four settings: no calling uncertainty, ancestry calling uncertainty only, genotype calling uncertainty only, and both ancestry and genotype calling uncertainty. Points denote replicate-averaged runtime, and intervals indicate 95% confidence intervals for runtime based on normal approximation.

**Figure S13:**
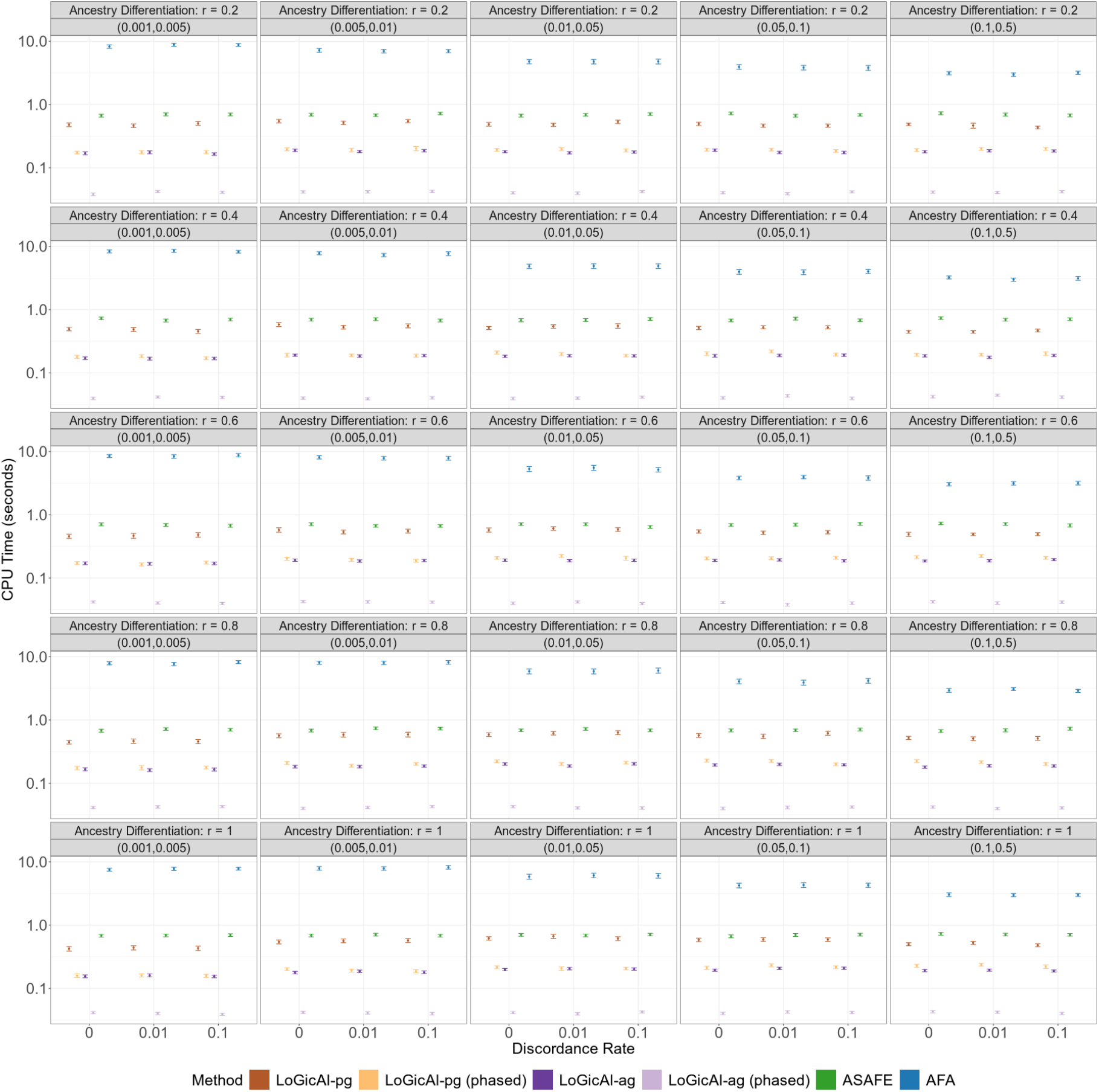
CPU runtime (seconds) of ancestry-specific allele frequency estimation methods across discordance rates, allele frequency bins and levels of ancestral population differentiation. We simulated 100, 000 phased individuals from a three-way admixture model under a scenario with probabilistic ancestry calling (*α* = 0.1) and a realistic genotyping error rate (*θ* = 5 × 10^−4^). Methods compared include LoGicAl (LoGicAl-pg, LoGicAl-pg (phased), LoGicAl-ag, LoGicAl-ag (phased)), ASAFE and AFA. Points denote replicate-averaged runtime, and intervals indicate 95% confidence intervals for runtime based on normal approximation.

**Figure S14:**
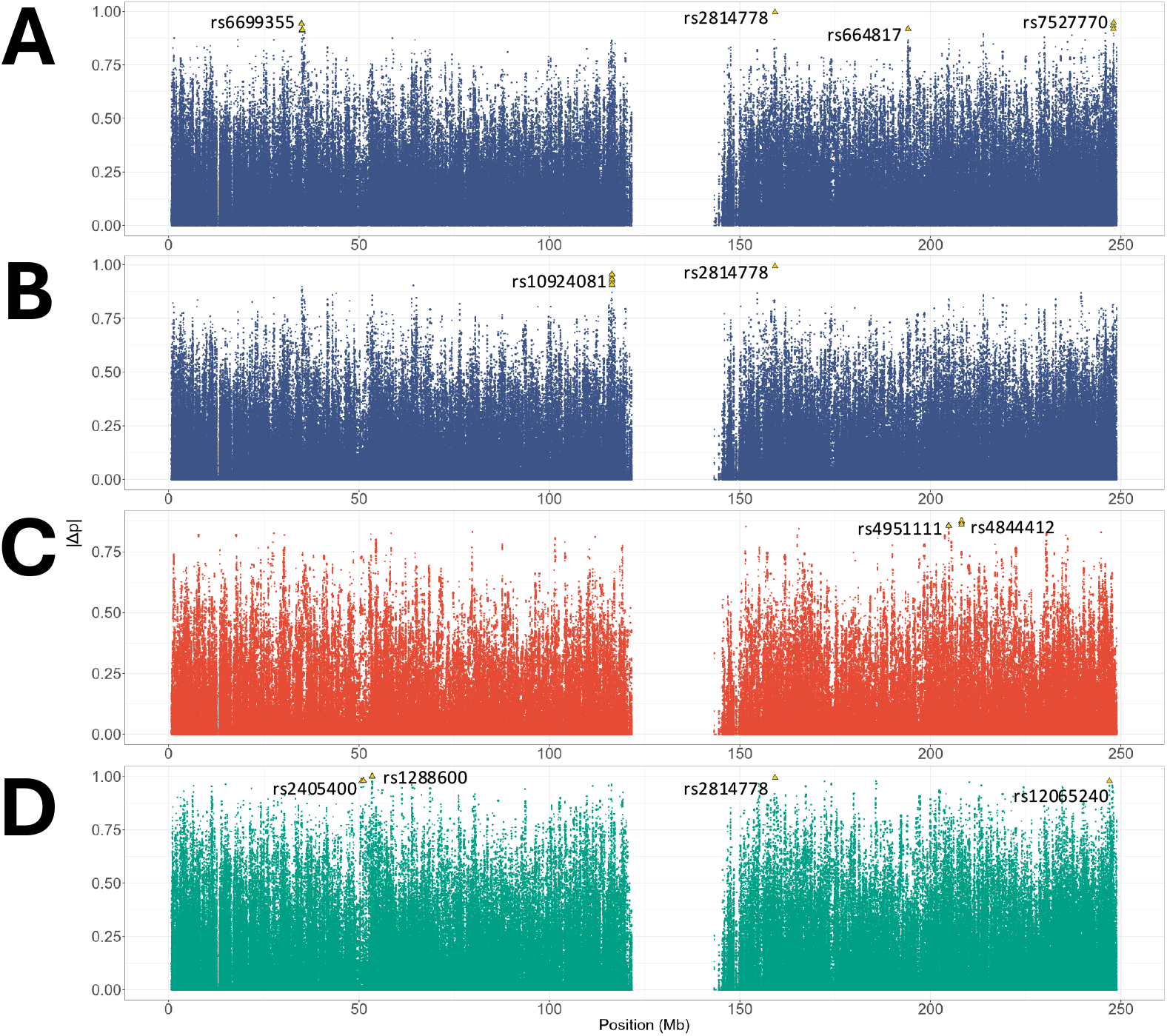
Absolute differences of ancestry-specific allele frequency estimates across chromosome 1. (A) Absolute differences between African and European ancestral estimates in 1kGP admixed African cohorts. (B) Absolute differences between African and European ancestral estimates in 1kGP admixed American cohorts. (C) Absolute differences between Native American and European ancestral estimates in 1kGP admixed American cohorts. (D) Absolute differences between Native American and African ancestral estimates in 1kGP admixed American cohorts. Outliers (yellow) denote the top 10 most differentiated variants per comparison; among these, the most differentiated variant within 1 Mb window is labeled by its rsID.

## Supplemental tables

**Table S1:**
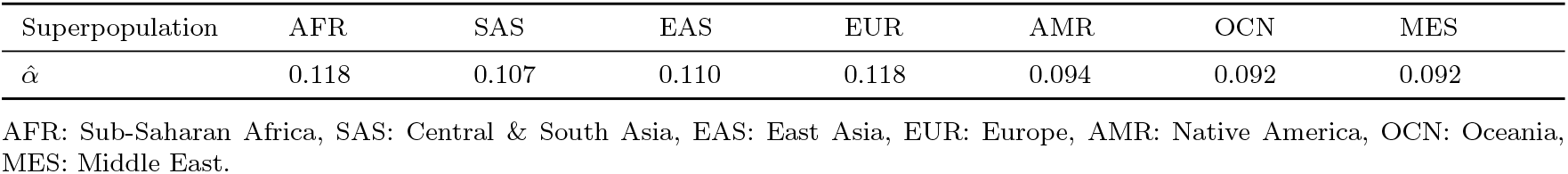
Dirichlet regression estimates of resolvability of local ancestry assignment across 1000 Genomes Project individuals. We fitted an intercept-only Dirichlet regression model to the posterior probabilities of assigning each ancestral superpopulation generated by RFMix, using the `DirichReg()`function from the `DirichletReg`R package. We calculated the estimated *α* parameters as the exponential of the coefficients corresponding to each superpopulation from the fitted model.

**Table S2:**
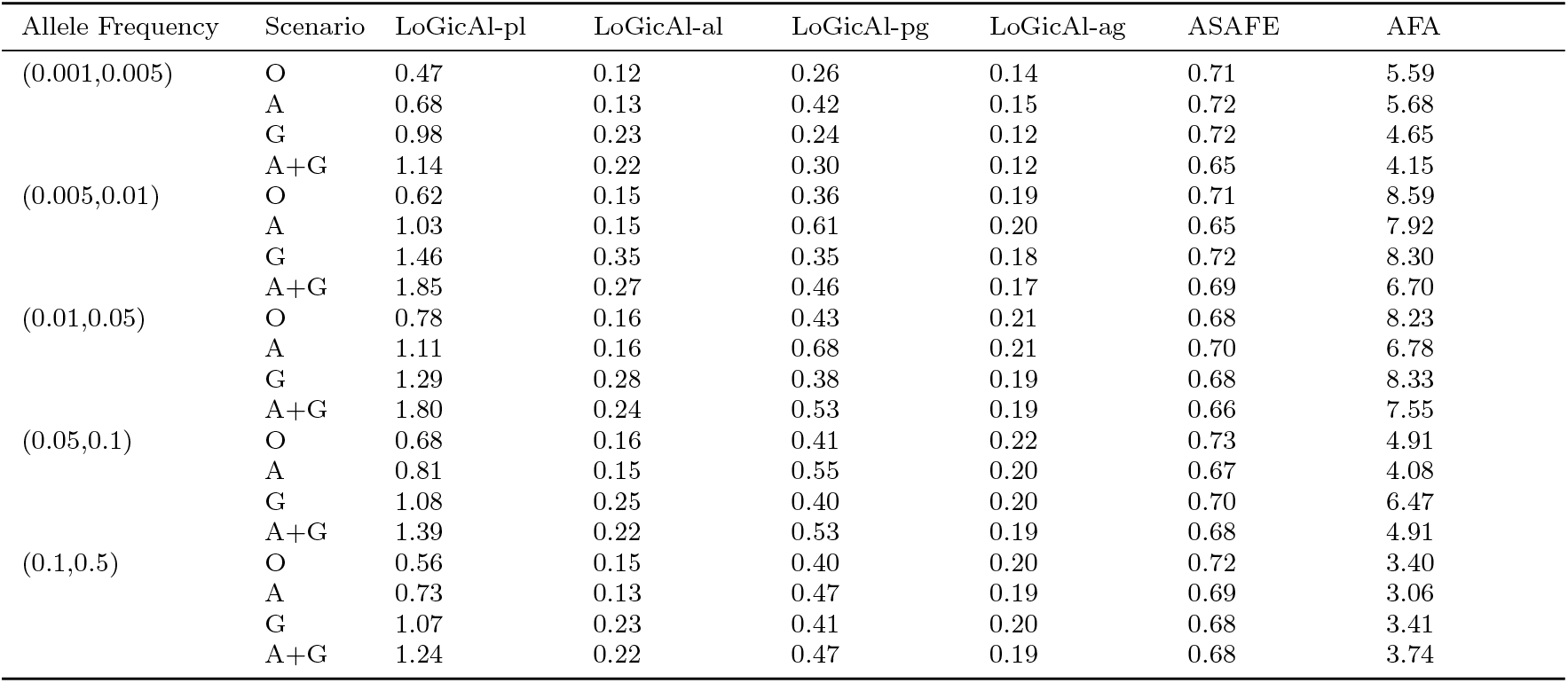
CPU runtime (seconds) of ancestry-specific allele frequency estimation methods across allele frequency bins and uncertainty scenarios. We simulated 100, 000 unphased individuals from a three-way admixture model with inter-continental differentiation (*r* = 1). Methods compared include LoGicAl (LoGicAl-pl, LoGicAl-al, LoGicAl-pg, LoGicAl-ag), ASAFE, and AFA under four settings: (O) no calling uncertainty, (A) ancestry calling uncertainty only, (G) genotype calling uncertainty only, and (A+G) both ancestry and genotype calling uncertainty. Runtimes are reported for a single variant and are averaged over 100 simulation replicates.

**Table S3:**
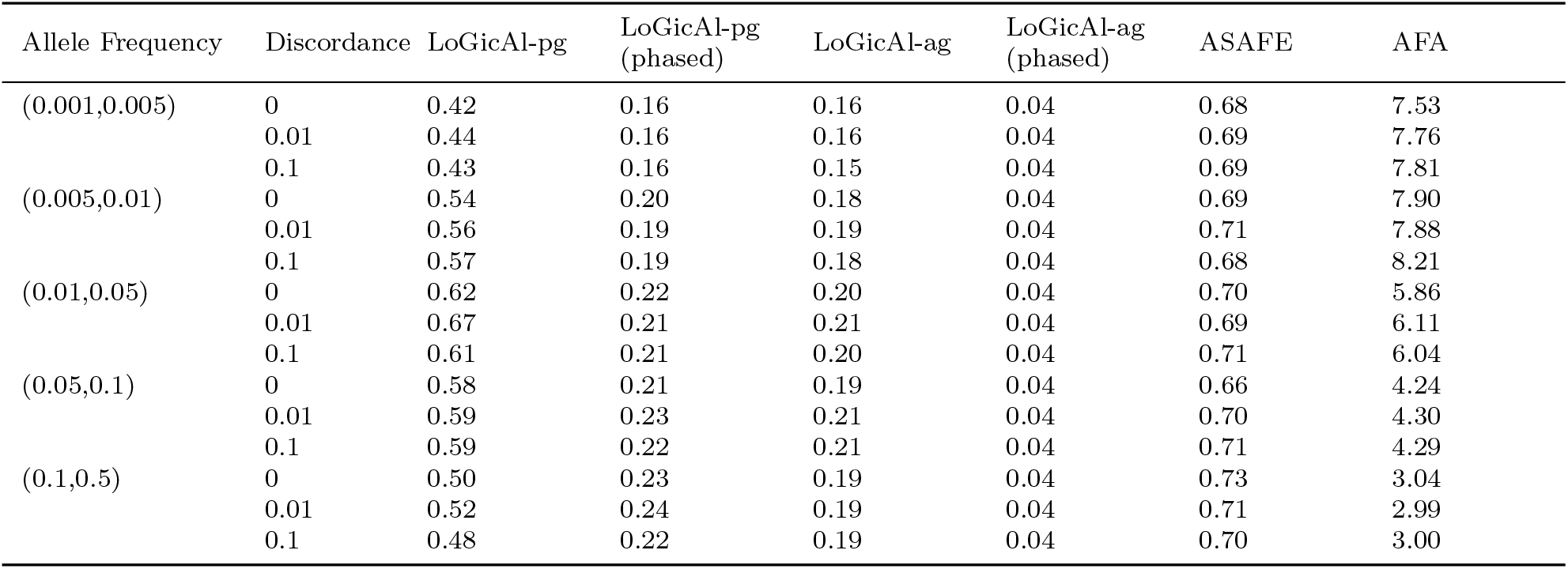
CPU runtime (seconds) of ancestry-specific allele frequency estimation methods across allele frequency bins and discordance rates. We simulated 100, 000 phased individuals from a three-way admixture model with inter-continental differentiation (*r* = 1). Methods compared include LoGicAl (LoGicAl-pg, LoGicAl-pg (phased), LoGicAl-ag, LoGicAl-ag (phased)), ASAFE, and AFA under a scenario with probabilistic ancestry calling (*α* = 0.1) and a realistic genotyping error rate (*θ* = 5 × 10^−4^). Runtimes are reported for a single variant and are averaged over 100 simulation replicates.

| Allele Frequency | Discordance | LoGicAl-pg | LoGicAl-pg (phased) | LoGicAl-ag | LoGicAl-ag (phased) | ASAFE | AFA |
| --- | --- | --- | --- | --- | --- | --- | --- |
| (0.001,0.005) | 0 | 0.42 | 0.16 | 0.16 | 0.04 | 0.68 | 7.53 |
|  | 0.01 | 0.44 | 0.16 | 0.16 | 0.04 | 0.69 | 7.76 |
|  | 0.1 | 0.43 | 0.16 | 0.15 | 0.04 | 0.69 | 7.81 |
| (0.005,0.01) | 0 | 0.54 | 0.20 | 0.18 | 0.04 | 0.69 | 7.90 |
|  | 0.01 | 0.56 | 0.19 | 0.19 | 0.04 | 0.71 | 7.88 |
|  | 0.1 | 0.57 | 0.19 | 0.18 | 0.04 | 0.68 | 8.21 |
| (0.01,0.05) | 0 | 0.62 | 0.22 | 0.20 | 0.04 | 0.70 | 5.86 |
|  | 0.01 | 0.67 | 0.21 | 0.21 | 0.04 | 0.69 | 6.11 |
|  | 0.1 | 0.61 | 0.21 | 0.20 | 0.04 | 0.71 | 6.04 |
| (0.05,0.1) | 0 | 0.58 | 0.21 | 0.19 | 0.04 | 0.66 | 4.24 |
|  | 0.01 | 0.59 | 0.23 | 0.21 | 0.04 | 0.70 | 4.30 |
|  | 0.1 | 0.59 | 0.22 | 0.21 | 0.04 | 0.71 | 4.29 |
| (0.1,0.5) | 0 | 0.50 | 0.23 | 0.19 | 0.04 | 0.73 | 3.04 |
|  | 0.01 | 0.52 | 0.24 | 0.19 | 0.04 | 0.71 | 2.99 |
|  | 0.1 | 0.48 | 0.22 | 0.19 | 0.04 | 0.70 | 3.00 |

**Table S4:** Allele frequency estimates for LoGicAl-identified strongest ancestry-differentiated variants on chromosome 1 in 1kGP admixed American cohorts.

| rsID | Gene Region | LoGicAl |  |  | gnomAD |  |  | Ancestry Pair |
| --- | --- | --- | --- | --- | --- | --- | --- | --- |
|  |  | AFR | EUR | NA | AFR | EUR | AMR |  |
| *rs2814778* | *ACKR1* | > 0.999 | 0.007 | 0.005 | 0.822 | 0.004 | 0.090 | AFR-EUR |
| *rs4844412* | *PLXNA2* | 0.221 | 0.108 | 0.985 | 0.145 | 0.151 | 0.378 | NA-EUR |
| *rs1288600* | *SLC25A3P1* | > 0.999 | 0.314 | < 0.001 | 0.858 | 0.273 | 0.275 | NA-AFR |

AFR: Sub-Saharan Africa, EUR: Europe, NA: Native America, AMR: Admixed America.

## Supplemental notes

### Note S1: Derivation of LoGicAl algorithm

Following the notations in section (2.2), the complete-data log-likelihood of ancestry-specific allele frequencies is

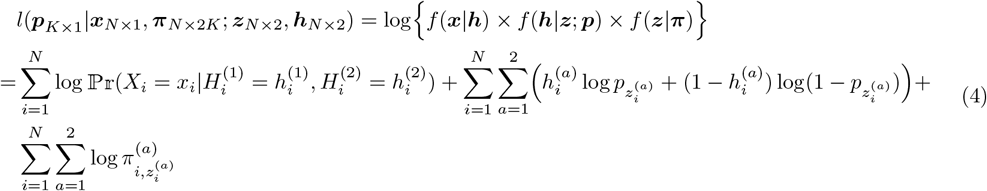

The joint conditional distribution of latent variables 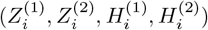 given the observed data and ancestry-specific allele frequencies is

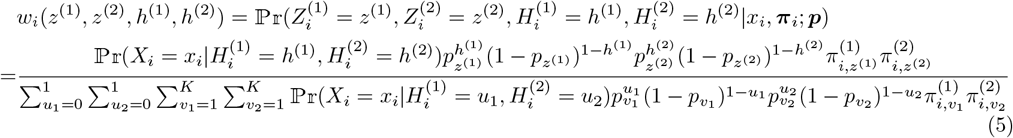

Therefore, the expected log-likelihood of ancestry-specific allele frequencies is

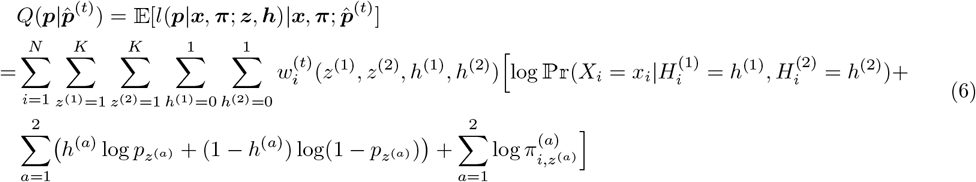

Taking derivative for each element *p*_*k*_ in ***p***

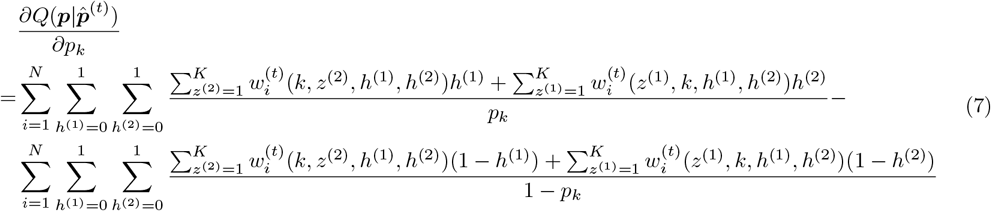

By setting 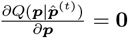, the expected log-likelihood 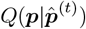 is maximized at

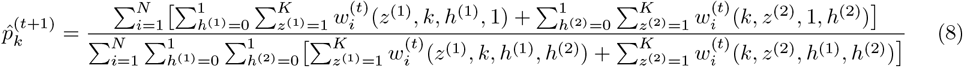

### Note S2: Simulation study details

We simulated ancestry-specific allele frequencies following the Balding-Nichols model[32] assuming each ancestral population descends independently from a common reference population. Under this model, the allele frequency *p*_*k*_ of the ancestral population *k* with reference allele frequency *p*_*A*_ is

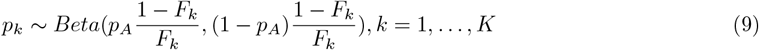

where *F*_*k*_ is Wright’s F-statistic[48, 49] measuring the genetic differentiation between the reference population and ancestral population *k*. Specifically, we simulated reference allele frequency *p*_*A*_ from a uniform distribution *p*_*A*_ ~ *U* (*a, b*) with varying (*a, b*) from (0.001, 0.005) to (0.1, 0.5) to generate allele frequency spectrum from rare to common variants. Based on the simulated *p*_*A*_, we sampled true ancestry-specific allele frequencies *p*_*k*_ from the distribution (9). Here, to generate ancestral population differentiation, we adopted the design of [19]: we first fixed the total number of ancestral populations at *K* = 3 and set (*F*_1_, *F*_2_, *F*_3_) = (0.01*r*, 0.05*r*, 0.1*r*), where *r* ∈ {0.2, 0.4, 0.6, 0.8, 1} is a scaling parameter controlling the degrees of population differentiation. This approach has been previously evaluated by [33, 50]. Larger *r* describes more rapid genetic drift rate from the reference population, making the descending populations more distinct from each other: setting *r* = 1 results in allele frequency differentiation observed between populations across different continents (e.g., between San, Han, and Maya populations), whereas setting *r* = 0.2 corresponds to differentiation between populations on the same continent[33]. Additionally, we varied the number of ancestral populations from *K* = 2 to *K* = 5 and set *F*_*k*_ = *F, k* = 1, …, *K*, with *F* = 0.01 to simulate weak population differentiation and *F* = 0.05 to simulate strong population differentiation.

To simulate true underlying local ancestries and their assignment probabilities, we sampled the assignment probabilities of ancestral groups for each individual independently from a Dirichlet distribution ***π***_*i*_ ~*Dir*(*α*𝟙_*K*_), where *α* quantifies the resolvability of local ancestry inference and we fixed *α* = 0.1 as estimated from the 1000 Genomes Project cohort (Table S1). Given the simulated assignment probabilities, we generated true local ancestry of each allele *a* from a multinoulli distribution 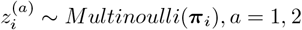, while generating the best-guess local ancestry call as the ancestral group where the assignment probability is maximized.

After obtaining the simulated ancestry-specific allele frequencies **p** and true local ancestries **z**^(**a**)^, *a* = 1, 2, we sampled true genotype per chromosome for each individual independently from a Bernoulli distribution 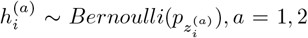. To imitate the effects of genotyping errors and statistical phasing uncertainty, we generated two types of observed genotype inputs: (1) genotype likelihoods representing probabilistic genotype information, and (2) best-guess genotype calls representing discreet genotype assignments.

We generated genotype likelihoods by adopting the framework in [51]. We simulated sequencing coverage for individual *i* from a Poisson distribution *d*_*i*_ ~ *Poisson*(*λ*_*i*_) after simulating average sequencing coverage from a truncated normal distribution *λ*_*i*_ ~ *TN* (*µ, µ/*3), *λ*_*i*_ ∈ (0, 2*µ*). Here, *µ* is average sequencing coverage across individuals and we chose *µ* ∈ {1, 5, 30} to model low-coverage to high-coverage sequencing data. We then sampled the number of sequencing reads presenting the alternate allele from a Binomial distribution 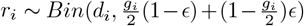, where 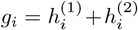 and *ϵ* is per-base sequencing error rate. We fixed *ϵ* = 0.001 across all simulation settings. Therefore, the genotype likelihoods for reference homozygote, heterozygote, and alternate homozygote are 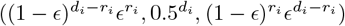, respectively. We used simulated overall allele frequency 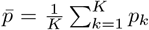 as prior and called genotypes for each individual as maximum-a-posteriori estimates based on the posterior probabilities 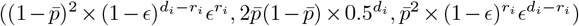. In parallel, to simulate genotype calls as observed genotype inputs, we generated observed genotype per chromosome from a Bernoulli distribution 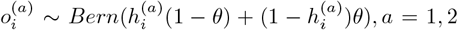, where *θ* is genotyping error rate and we fixed *θ* = 5 × 10^−4^ as estimated from previous literature[38] to mimic error rates from either array-based (e.g., UK Biobank[52]) or sequence-based (e.g., TOPMed[45]) genotype calling. To simulate observed phased genotypes, we exchanged the values of 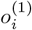 and 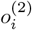 with probability *γ*, generating observed phase 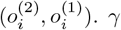 is a compound discordance rate measuring deviations of observed ancestry-genotype phase from true ancestry-genotype phase. These deviations can arise from switch errors and double switch errors (i.e., flip errors) in statistical phasing[38] and inferred ancestry segment flips[39]. Following reported values[44], we varied *γ* ∈ {0, 1%, 10%}.

